# Anthracyclines induce global changes in cardiomyocyte chromatin accessibility that overlap with cardiovascular disease loci

**DOI:** 10.1101/2025.06.11.658997

**Authors:** E. Renee Matthews, Raodatullah O. Abodunrin, John D. Hurley, José Angel Gutiérrez, Michelle C. Ward

## Abstract

Breast cancer drugs including anthracyclines (ACs) and Trastuzumab increase the risk for cardiovascular diseases (CVDs) such as atrial fibrillation (AF) and heart failure (HF) that ultimately affect the heart muscle. These CVDs are associated with hundreds of genetic variants in non-coding regions of the genome. However, how these drugs affect the regulatory potential of the non-coding genome of the heart and CVD risk loci is unknown. We therefore measured global chromatin accessibility across iPSC-derived cardiomyocytes from four individuals treated with the ACs, Doxorubicin, Epirubicin, and Daunorubicin, a related non-AC, Mitoxantrone, and the monoclonal antibody Trastuzumab, or a vehicle control for three and 24 hours. We identified 155,557 high-confidence regions of open chromatin across 48 samples where the major sources of variation are associated with drug type and time. Jointly modeling the data revealed three accessibility response signatures denoted as early-acute, early-sustained, and late that correspond to 67,329 regions that open or close in response to drug treatment. Sequences associated with drug-induced chromatin opening contain motifs for DNA damage-associated transcription factors including p53 and ZBTB14, and associate with increases in active histone acetylation and gene expression. 21 AF- and HF-associated SNPs directly overlap with regions associated with drug-induced opening. A shared intronic HF and AF SNP, rs3176326, that is also an eQTL for *CDKN1A* in heart tissue, associates with increased chromatin accessibility, histone acetylation, and *CDKN1A* expression in response to all ACs. Our results demonstrate large-scale changes in chromatin accessibility in cardiomyocytes treated with ACs, which correspond to several regions harboring CVD risk loci.

**Author summary:** Anthracyclines are a widely used class of breast cancer drugs that are linked to cardiac toxicity and the development of heart disease in some women. There are hundreds of genetic variants that associate with risk for heart disease; however their role and mechanism of action in drug-induced toxicity is unclear given that most reside in the non-coding genome. We therefore tested the effects of five breast cancer drugs on genome-wide chromatin accessibility using induced pluripotent stem cell-derived cardiomyocytes. We found tens of thousands of chromatin regions that change in accessibility after drug treatment. Regions with increased accessibility contain sequences that have been shown to bind DNA damage-associated transcription factors, and associate with increases in nearby gene expression. We find 21 heart disease-associated genetic variants in regions that increase in chromatin accessibility following treatment. This suggests that cancer drugs have large effects on the non-coding genome of heart cells including at regions associated with heart disease. This research contributes to our understanding of how genetic variants associated with disease exert their effects.

## Introduction

Anthracyclines (ACs) and the monoclonal antibody Trastuzumab (TRZ) are effective therapeutic agents used in the treatment of breast cancer. TRZ is widely used to treat HER2-positive breast cancers, while ACs including Doxorubicin (DOX) and its structural analogues Epirubicin (EPI) and Daunorubicin (DNR), are standard treatments for triple-negative breast cancers in addition to being used to treat other cancer types regardless of HER2 receptor status.

Treatment with either ACs or TRZ associates with increased risk for cardiovascular disease (CVD) and can ultimately lead to heart failure (1, 2). Epidemiological studies have indicated that breast cancer patients treated with ACs are more likely to develop heart failure than healthy individuals (3). Even young breast cancer patients, without CVD risk factors, show decreased heart contraction following AC treatment (4). In addition to heart failure, AC treatment also associates with increased risk for arrhythmias (5). Notably the increased risk for CVD in breast cancer patients is associated with heart failure and arrhythmia, and not ischemic heart disease, indicating that damage to the myocardium itself is evident (6).

Cardiac dysfunction following cancer treatment can be referred to as cancer therapy-related cardiac dysfunction (CTRCD)(7). While the exact definition varies across clinical guidelines, declining heart function, typically deduced through a reduction of the heart’s ejection fraction, is accepted as a key metric. To understand the genetic component of risk for breast cancer patients developing CTRCDs, Genome-wide association studies (GWAS) have been performed (8–10). These studies have highlighted tens of genetic variants nominally-associated with AC toxicity (8, 9). Over a hundred genetic variants have also been robustly associated with heart failure and arrhythmia independent of cancer drug treatment, indicating a genetic component to CVD (11).

Deducing the mechanisms behind, and effects of, disease-associated genetic variants is challenging in human patients. Large numbers of individuals are required to overcome environmental effects, and controlled perturbation studies cannot typically be performed for ethical and technical reasons. The ability to generate disease-relevant cell types from induced pluripotent stem cells has facilitated progress in this area. Cellular models using induced pluripotent stem cell-derived cardiomyocytes (iPSC-CMs) have recapitulated the clinically-observed AC- and TRZ-induced cardiotoxicity phenotype, validating this *in vitro* approach (12, 13). These cellular models have allowed for the identification of genetic variants that associate with gene expression in response to DOX providing further support for the role of genetic variation and its molecular effects on cardiotoxicity (14). We have shown that treatment with ACs (DOX, DNR, EPI) impacts the transcriptome in a similar manner across drugs and affects cardiomyocyte function (15). However, the molecular basis for the gene expression changes, and how they contribute to cardiotoxicity is unclear.

ACs have classically been characterized as DNA-damaging agents that induce DNA double strand breaks through their interactions with the topoisomerase TOP2 (16, 17). However, there is more recent evidence that ACs induce their effects through damage to both DNA and chromatin, and that decoupling these effects can limit cardiotoxicity (18). Survival of breast cancer patients after DOX treatment has been shown to be mediated through chromatin regulators (19). Chromatin regulators are also enriched amongst genes that respond to ACs in iPSC-CMs implicating a chromatin mechanism in cardiomyocytes (15). DOX has also been shown to affect chromatin accessibility around transcription factor motifs in breast cancer cells suggesting a mechanism for transcriptional changes (20). However, the impact of ACs and TRZ on the chromatin landscape of cardiomyocytes is unknown.

We therefore designed a study to investigate the effects of five breast cancer drugs including three ACs, the anthracenedione Mitoxantrone (MTX) that also targets TOP2, and TRZ on the global chromatin landscape of iPSC-CMs in four healthy female individuals. We were able to identify thousands of genomic regions where chromatin accessibility changes in response to drug treatment, and investigate the chromatin context of CVD-associated genetic variants in the presence of cancer drug treatment.

## Results

### AC treatment induces thousands of chromatin accessibility changes in iPSC-CMs

To test the effects of breast cancer drugs on cardiomyocytes, we obtained iPSCs from four healthy female individuals and differentiated them into iPSC-CMs. The median proportion of cells expressing cardiac troponin T across individuals is 98% indicating high-purity cultures (S1 Table). A previous study has indicated that treatment of iPSC-CMs with a sub-lethal, clinically-relevant concentration (0.5 µM) of the TOP2 inhibitors (TOP2i) DOX, DNR, EPI and MTX induces thousands of gene expression changes, but TRZ does not (15). To determine whether these DNA-damaging agents also have effects on chromatin accessibility, we treated iPSC-CMs from four individuals with DOX, DNR, EPI, MTX, TRZ and a vehicle control (VEH) and collected cells for global chromatin analysis by ATAC-seq at three and 24 hours post-treatment in treatment-balanced batches (Fig 1A & S1 Table).

**Figure 1:**
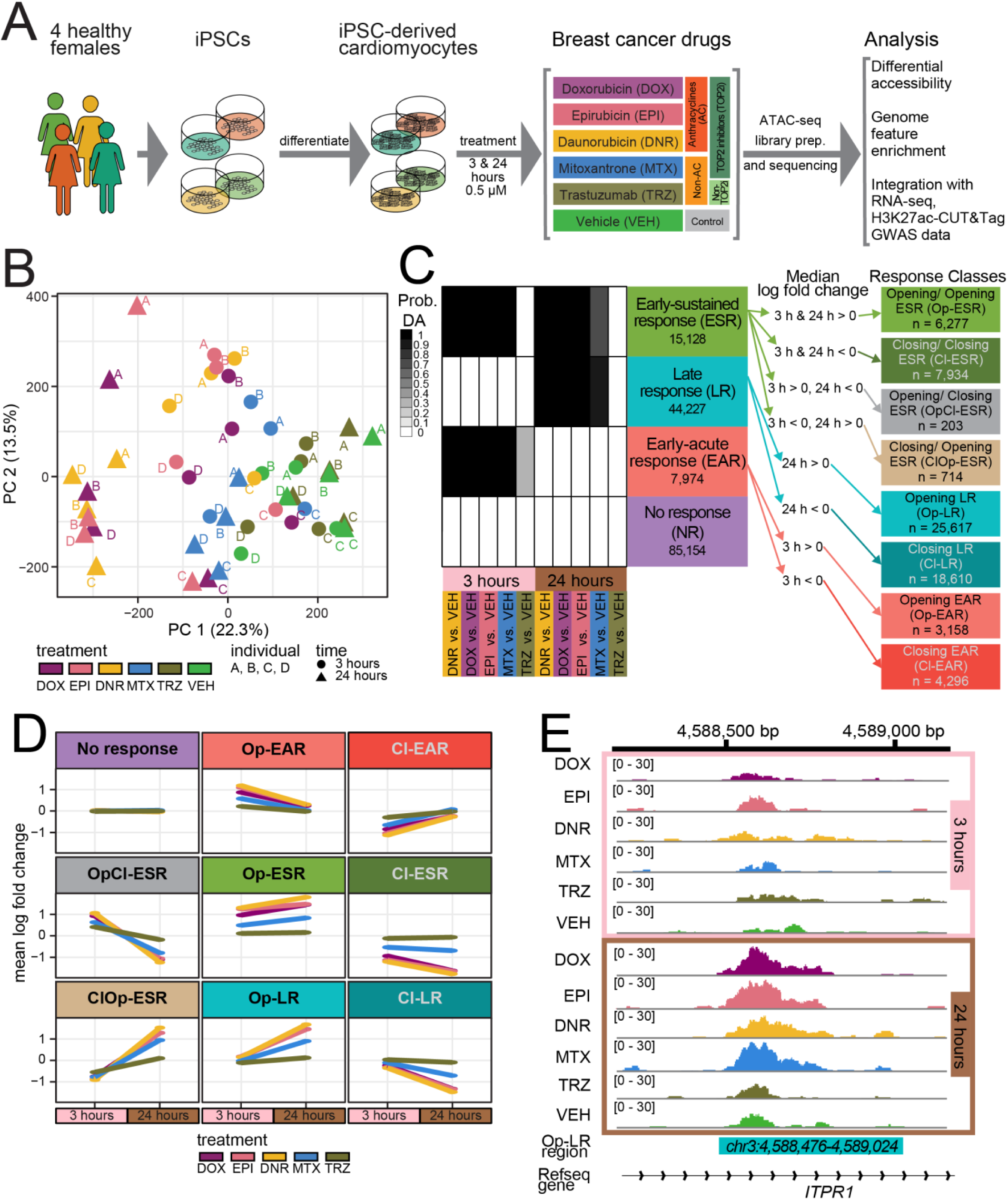
Thousands of chromatin accessibility changes are induced in iPSC-CMs following anthracycline treatment. **(A)** Experimental design of the study. iPSCs derived from four healthy women aged 20 to 30 were differentiated into cardiomyocytes (iPSC-CMs) and exposed to a panel of drugs used in breast cancer treatment. Chromatin accessibility changes in response to TOP2i Doxorubicin (DOX), Epirubicin (EPI), Daunorubicin (DNR), and Mitoxantrone (MTX), a non-TOP2i, Trastuzumab (TRZ), and a water vehicle (VEH) were measured by ATAC-seq. **(B)** Principal component analysis of ATAC-seq-derived accessibility measurements (log_2_ cpm) across 48 samples representing four individuals (A,B,C,D), the length of exposure (three hours: circles, 24 hours: triangles), and six treatments (DOX (mauve), EPI (pink), DNR (yellow), MTX (blue), TRZ (olive) and VEH (green). **(C)** Chromatin accessibility response classes identified following joint Bayesian modeling of test pairs. Shades of black represent posterior probabilities of chromatin being differentially accessible in response to each drug treatment compared to VEH at each timepoint. Regions are categorized into four classes based on their posterior probabilities: ‘Early-acute response’ (EAR: red), ‘Early-sustained response’ (ESR: green), ‘Late response’ (LR: blue), ‘No response’ (NR: purple). Response classes are further separated into opening or closing chromatin regions in response to treatment based on the median log fold change across drugs at each timepoint based on a pairwise differential accessibility analysis comparing each drug to the VEH. This results in nine response classes: NR, opening EAR (Op-EAR), closing EAR (Cl-EAR), opening ESR at both timepoints (Op-ESR), closing at both timepoints (Cl-ESR), early opening late closing (OpCl-ESR), early closing and late opening (ClOp-ESR), opening LR (Op-LR), closing LR (Cl-LR). **(D)** Mean log fold change of open chromatin regions across drug treatments compared with VEH at each timepoint for regions in each of the nine response classes. Treatments are shown with the following colors: DOX (mauve), EPI (pink), DNR (yellow), MTX (blue), and TRZ (olive). **(E)** Example of an accessible chromatin region within an intron of the *ITPR1* gene, classified as Op-LR, across drug treatments and time for a representative individual (Individual D).

For each of the 48 samples, we obtained ATAC-seq reads that map uniquely to the nuclear genome, and designated open chromatin regions (See Methods; Fig S1 & S2 Table). The number of mapped fragments and open chromatin regions is similar across treatments within each timepoint (Fig S2). The median number of open chromatin regions is higher in the samples treated for 24 hours than those treated for three hours (3 hour median = 73,241, 24 hour median = 88,737). In order to identify a high-confidence set of open chromatin regions we removed annotated blacklist regions that are known to exhibit non-specific signal, and selected those regions that are present in at least five of the 48 samples. This yielded a superset of 172,481 open chromatin regions. To assess the quality of the ATAC-seq data we determined the percentage of fragments in the set of open chromatin regions for each sample. All samples have at least 20% of fragments in open chromatin regions in line with standards of the field (Fig S3).

In order to quantify changes in chromatin accessibility following treatment, we counted the number of reads in the superset of open chromatin regions across all samples. We filtered out regions with low read counts yielding 155,557 regions for downstream analysis. To determine whether the accessible chromatin regions we identified are physiologically relevant, we compared our data to chromatin accessibility data from heart ventricle tissue from a female donor (21). We found that 43% of our accessible regions overlap with regions identified in heart tissue (Fig S4). These open chromatin regions are also enriched at transcription start sites (TSS; Fig S5A-B), including that of *TNNT2*, a cardiac-specific gene (Fig S6). Unsupervised hierarchical clustering separates samples based on whether they are treated with TOP2i for 24 hours, and then by time and treatment group (Fig S7). Principal component analysis revealed that the first principal component, representing 22.31% of variation in the data associates with treatment, and the second principal component, representing 13.53% of variation in the data, associates with individual and time (Fig 1B, Fig S8A-C).

We next tested the effect of each drug on chromatin accessibility at each timepoint. We identified thousands of differentially accessible regions (DARs; adjusted *P* < 0.05) following three hours of TOP2i treatment (DOX vs VEH = 3,473; DNR vs VEH = 22,738; EPI vs VEH = 14,234; MTX vs VEH = 804; TRZ vs VEH = 1; Fig S9 and S3-7 Tables), and tens of thousands after 24 hours of treatment (DOX vs VEH = 64,820; DNR vs VEH = 79,995; EPI vs VEH = 66,501; MTX vs VEH = 24,250; TRZ vs VEH = 0; Fig S9 and S8-12 Tables). There was only one DAR following TRZ treatment at three hours and no DARs following TRZ treatment at 24 hours. The most significant DAR following each AC treatment is also a DAR in the other AC treatments and includes loci near the heat shock protein gene *DNAJB12* and the replication gene *FBXL12* (Fig S10A). Similarly, log fold changes across ACs are highly correlated at each timepoint (Fig S10B) suggesting sharing of the chromatin response across ACs.

In order to determine the changes in chromatin accessibility across drugs and time we used a joint Bayesian model. We first determined the number of clusters that explain the predominant accessibility patterns in the data using Bayesian Information Criterion and Akaike Information Criterion analysis. This analysis revealed four sets of regions (Fig S11). Each cluster is driven by timepoint and not drug type. We therefore labelled the four categories as No Response regions (NR; n = 85,154), Early Acute Response regions (EAR, n = 7,974), Early Sustained Response regions (ESR, n = 15,128), and Late Response regions (LR, n = 44,227; Fig 1C). These response categories identify shared patterns of accessibility changes irrespective of the direction of the effect. We further categorized the four sets of response regions into those that are opening or closing in response to drug treatment. We used the log fold change information for each drug from the pairwise differential accessibility analysis, calculated the median fold change across drugs at each timepoint, and used this value to separate regions in each response category into those that are opening and closing (Fig 1C). This resulted in 4,899 opening EAR regions (Op- EAR; median log fold change > 0 at 3 hours), 3,075 closing EAR regions (Cl-EAR; median log fold change < 0 at 3 hours), 25,617 opening LR (Op-LR; median log fold change > 0 at 24 hours), and 18,610 closing LR (Cl-LR; median log fold change < 0 at 24 hours). We categorized the ESR regions into those that are open at three and 24 hours (Op-ESR, n= 6,277), regions that are closed at both timepoints (Cl-ESR, n = 7,934), and regions that show the opposite direction at each timepoint (OpCl-ESR, n = 203 and ClOp-ESR, n = 714). Categories of regions show the expected accessibility profile across treatments and time (Fig 1D-E). We used these nine categories of regions moving forward (S13 Table).

### Early drug-responsive chromatin regions are enriched in CpG-rich regions near transcription start sites

We were next interested in identifying sequence features associated with our set of nine chromatin response regions. Given that approximately half of the human genome comprises transposable elements (TEs), we first calculated the proportion of TEs overlapping each chromatin category and calculated enrichment in each response category relative to the non-response category. The Op-LR is the only category enriched for all TEs (Chi-square test; adjusted *P* < 0.05; Fig 2A). A TE is said to overlap a chromatin region when it overlaps by 1 bp. Results are similar when requiring a chromatin region to overlap 50% of the total length of a TE. We next considered each class of TE independently. The Op-LR category is enriched for most TE classes considered including SINE, LINE, LTR, and DNA transposon. However, the TE class with the most notable enrichment is the SVA retrotransposon class that is highly enriched in both the Op- EAR and Op-ESR categories. We therefore investigated the constituent TEs within this class across the response categories and observed a strong contribution from SVA_D elements (Fig 2B). SVA elements have previously been shown to be co-opted into regulatory elements (22). Given that SVA TEs are CpG-rich elements, we next asked whether these response categories are enriched in CpG-dense CpG islands. We find the only enriched category for CpG islands is Op-EAR (Fig 2A). As most CpG islands are located close to transcription start sites (TSS), we tested for enrichment of this feature across our response categories. Only the Op-EAR category is enriched at TSS (Fig 2A).

**Figure 2:**
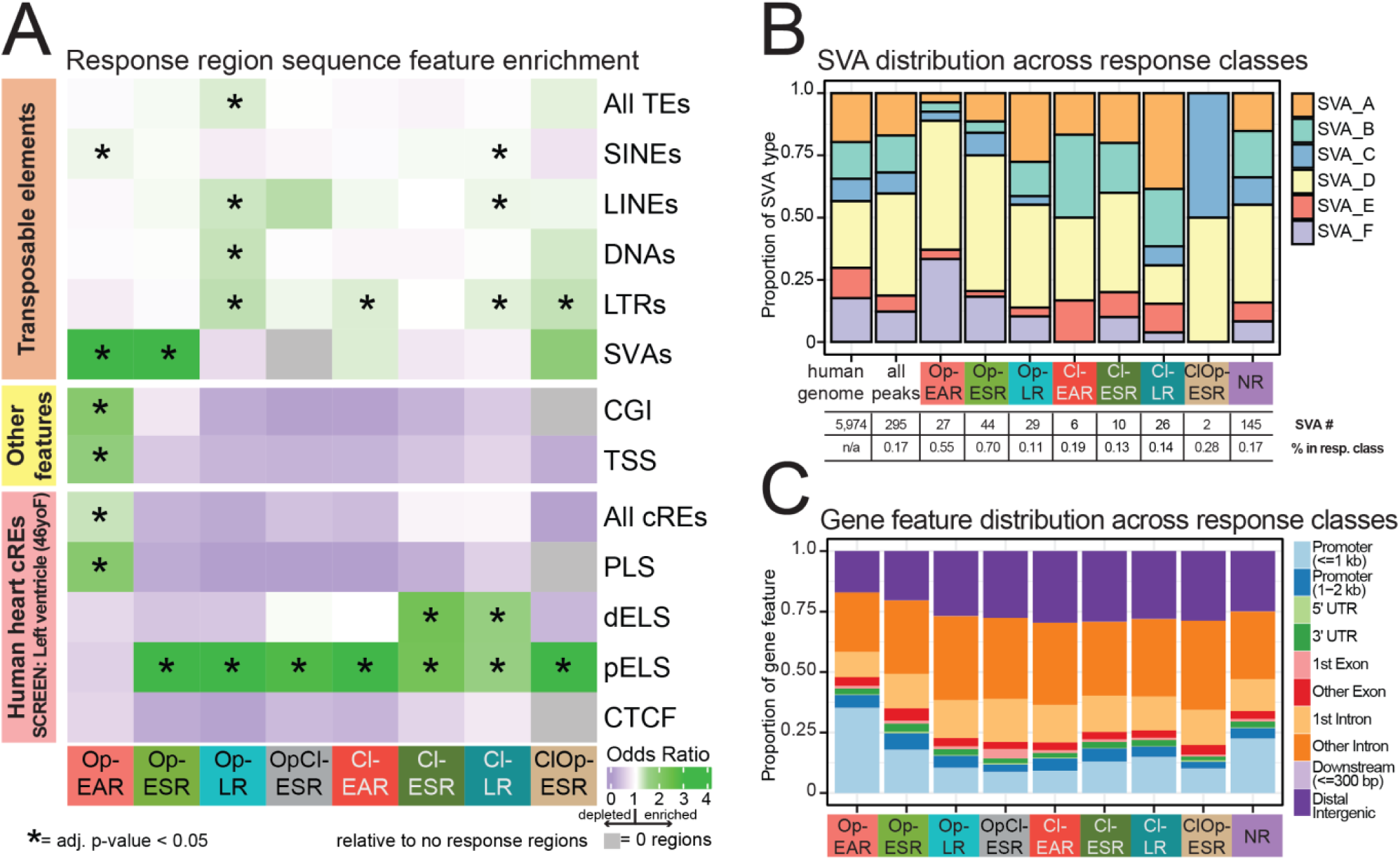
Early drug-dependent responsive chromatin regions are enriched in CpG-rich regions near transcription start sites. **(A)** Enrichment of genomic features in chromatin response classes compared to the No response class determined by chi-square test and Benjamini-Hochberg multiple testing correction. Transposable elements (TE) are represented in aggregate, and stratified into five TE classes; short-interspersed nuclear elements (SINEs), long-interspersed nuclear elements (LINEs), DNA transposons (DNAs), long terminal repeats (LTRs), and SINE-VNTR-*Alu* retrotransposons (SVAs). The ‘Other features category’ consists of CpG islands (CGI) and transcription start sites (TSS). Human heart candidate *cis*-regulatory elements (cREs) from a female adult were obtained from the ENCODE SCREEN database (21). cREs are represented in aggregate (All cREs) and stratified into promoter-like sequences (PLS), distal enhancer-like sequences (dELS), proximal enhancer-like sequence (pELS), and CCCTC-binding factor sequences (CTCF). **(B)** Proportion of SVA element types in the human genome, all accessible chromatin, and across chromatin response classes. The number of SVA elements found in each response class is listed below the class name, along with the percentage of open chromatin regions that contain an SVA element within that class. **(C)** Distribution of gene features across regions in each response class. ‘Downstream’ refers to regions where the center of the region is located less than 300 bp from the transcription termination site. ‘Distal intergenic’ refers to regions not localized within any other gene feature.

We next asked how these chromatin regions correspond to regulatory elements previously identified from DNase I hypersensitivity data and ChIP-seq data for H3K27ac, H3K4me3 and CTCF in human heart tissue from the ENCODE SCREEN database (See Methods). We considered all defined *cis*-regulatory elements (cREs) and found only the Op-EAR category to be enriched (Fig 2A). This category is also the only category enriched for promoter-like sequences (PLS), compared to all other chromatin region sets that are enriched for proximal enhancer-like sequences (pELS) or distal enhancer-like sequences (dELS). When considering gene feature annotations, the Op-EAR category is enriched for promoter sequences within 1 kb of TSS unlike the other categories, which is in line with the results indicating the Op-EAR category is enriched at TSS and PLS (Fig 2C). Together, these results suggest that the set of chromatin regions that open early in response to drug treatment has distinct sequence features and genomic distribution compared to regions that open later or close in response to drug treatment.

### Regions that open in response to AC treatment contain motifs for transcription factors implicated in the DNA damage response

Given that open chromatin region categories are enriched for regulatory elements, we next asked whether there are sequence motifs for transcription factors enriched within our chromatin response regions compared to regions that do not change in accessibility in response to drug treatment. Regions that open in response to drugs are enriched for motifs of DNA damage responsive transcription factors such as ZNF93 and ZBTB14, (Op-EAR; Fig 3A & S14 Table). ZNF93 is a KRAB zinc-finger transcription factor that is associated with DNA damage survival response in drug-resistant chondrosarcoma cells (23), and directly silences SVA elements in the human genome (24). ZBTB14, also known as ZNF478 and ZFP161, is involved in responding to replication stress (25) and DNA damage (26). In regions that open early and remain open (Op-ESR), TP53 and GLI1 motifs are enriched (Fig 3B). The p53 transcription factor is known to be a key factor in the DOX-induced DNA damage response in cardiomyocytes (27). GLI1 has been shown to regulate DNA damage response genes (28). Late response open regions are enriched for POU4F1 and FOS::JUNB motifs. POU4F, also known as BRN3A, has been shown to bind with p53 at the *BCL-2* promotor to suppress its activation (29). Regions that open and then close contain motifs for TEAD3 and HOXC10 (OpCl-ESR; Fig 3D). TEAD3 associates with DNA damage repair proteins and is required for genome stability (30), while HOXC10 is linked to breast cancer drug resistance through facilitating DNA damage repair (31).

**Figure 3:**
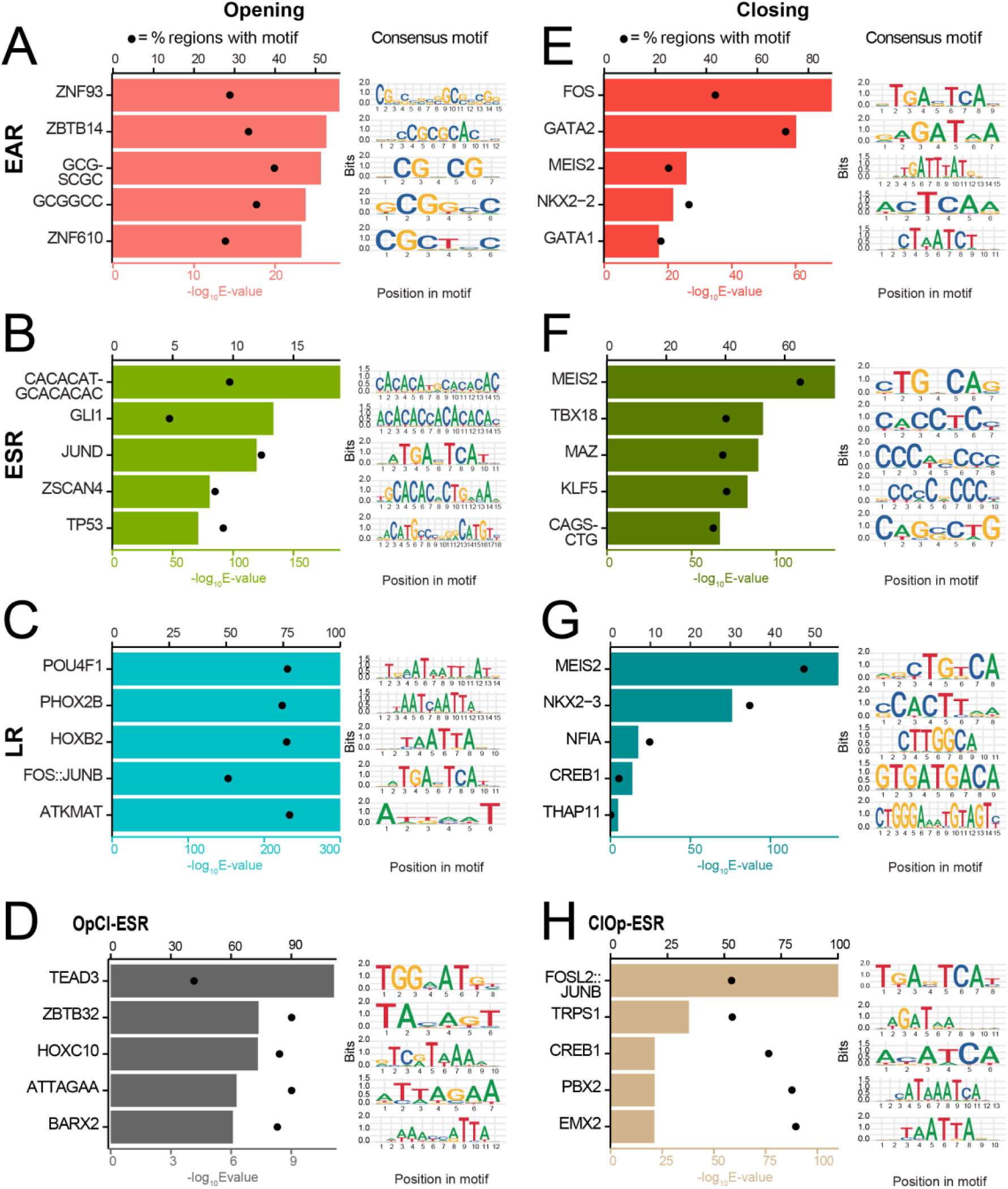
Regions opening in response to drug treatment are enriched for transcription factor motifs including TP53 and JUND. (A-H) Top five transcription factor consensus motifs enriched in Op-EAR (A), Op-ESR (B), Op-LR (C), OpCl-ESR (D), Cl-EAR (E), Cl-ESR (F), Cl-LR (G), or ClOp-ESR (H) regions compared to NR regions. Enriched motifs (– log_10_(E-value) < 0.05) with an enrichment ratio value > 1.25 compared to NR were selected. The top five consensus motifs unique to each motif cluster based on E-value are shown for each response class. Top x-axis displays percentage of regions in the class that have the consensus motif (data represented as circles). Bottom x-axis displays enrichment (– log_10_(E-value)) as calculated by XSTREME (data represented as bars). Motif logos for each enriched consensus motif are shown.

Regions that close in response to drugs are enriched for FOS and GATA2 (Cl-EAR; Fig 3E), MEIS2 and TBX18 (Cl-ESR; Fig 3F), MEIS2 and NFIA (Cl-LR; Fig 3G), and FOSL2::JUNB and TRPS1 (ClOp-ESR; Fig 3H).

### Regions opening in response to drugs associate with signaling pathways while those closing associate with cardiac-related pathways

To understand the impact of changing chromatin accessibility, we associated each of our chromatin regions in the nine different response categories with the TSS of the nearest gene expressed in iPSC-CMs (15), and asked which biological processes associate with these gene sets (Fig 4A). Regions that open in response to drugs are enriched (adjusted *P* < 0.05) for metabolic processes (Op-EAR), and developmental processes (Op-ESR), Fig 4B-C, S15 Table). Regions that close in response to drugs are enriched for signaling and development-related processes (Fig 4D-E, S15 Table), Cl-EAR is enriched for external stimulus response and animal organ development and Cl-ESR is enriched for organism development processes, heart contraction and heart processes. Regions classified as NR were not enriched in specific biological or cellular processes (Fig 4F). This suggests that transcriptional programs are either activated or repressed in response to opening and closing chromatin regions.

**Figure 4:**
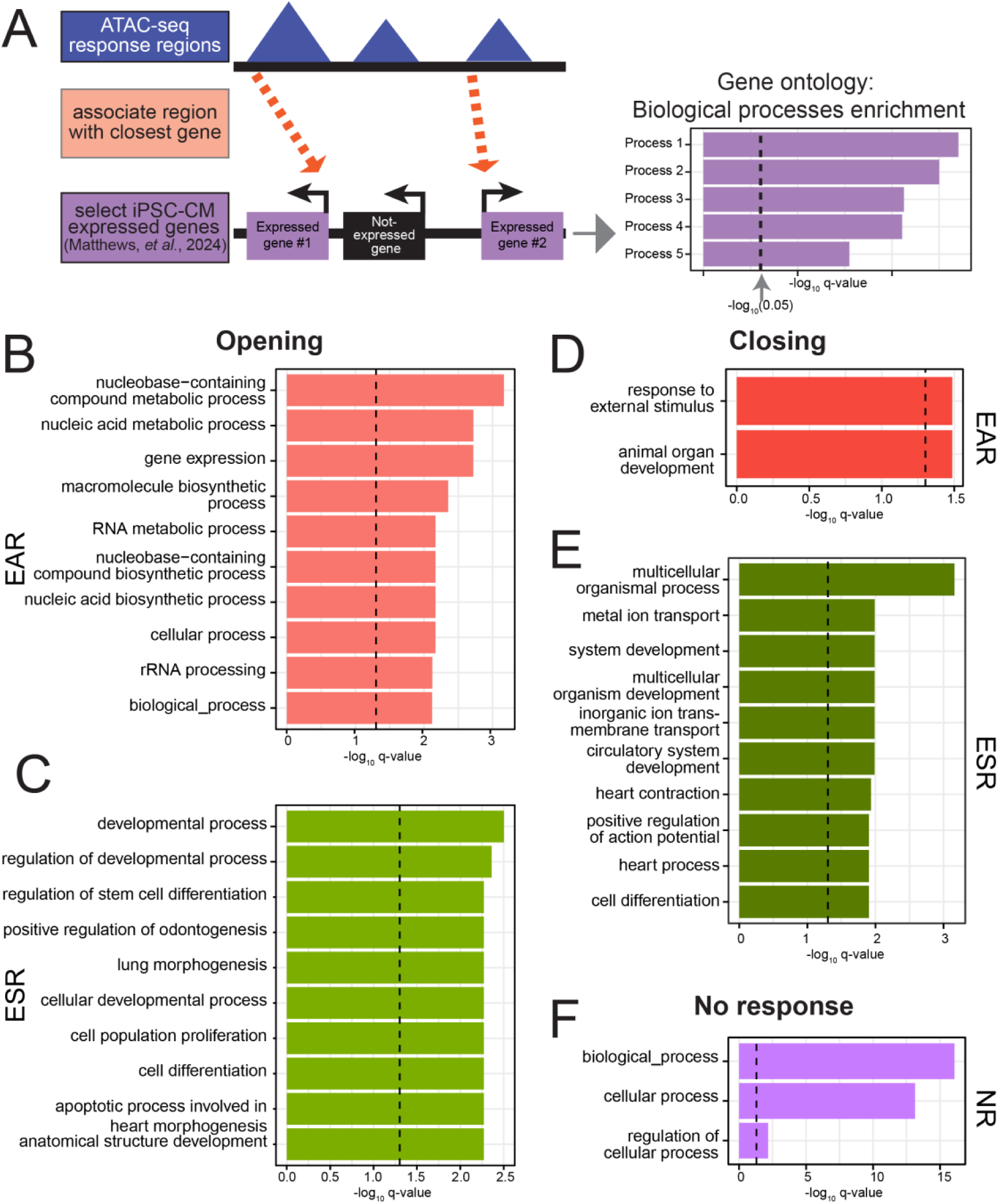
Regions that open in response to drugs are enriched near signaling genes, while closing regions are enriched near cardiac genes. **(A)** Schematic of the process to associate accessible chromatin response regions with genes expressed in iPSC-CMs. The TSS of the set of genes expressed in iPSC-CMs (15) was used to associate each response region with a gene. Unique genes from each response class were tested for biological process enrichment compared to the background of all expressed genes (n = 14,084). **(B-F)** Top 10 enriched biological processes (Benjamini-Hochberg q value < 0.05) for each class. The Cl-LR and Op-LR classes did not show enrichment for any processes. The dotted lines represent the –log_10_(q value) = 0.05.

### AC-induced chromatin accessibility changes correspond to changes in active histone modifications and gene expression

Chromatin regions that change in accessibility in response to drugs may coincide with gene regulatory regions. To test this, we selected a subset of individuals and profiled the enrichment of the active histone modification H3K27ac by CUT&Tag in TOP2i and VEH-treated iPSC-CMs. We used cells from the same differentiation and treatment batch used to generate the ATAC-seq data (S16 Table). We identified regions enriched for H3K27ac as we did for chromatin accessibility (See Methods; S17 Table). This yielded 20,137 regions enriched for H3K27ac in at least four samples. These regions are enriched around TSS as expected (Fig S12). 98.8% of H3K27ac regions overlap with our regions of accessible chromatin. Following removal of outlier samples, the samples primarily separate based on whether they were treated with TOP2i at 24 hours, similar to the chromatin accessibility data (Fig S13). We identified hundreds of differentially enriched regions (DERs; adjusted *P* < 0.05) following three hours of TOP2i treatment (DOX vs VEH = 114; DNR vs VEH = 1,277; EPI vs VEH = 0; MTX vs VEH = 0; Fig S14A; S18-21 Tables), and hundreds to thousands after 24 hours of treatment (DOX vs VEH = 840; DNR vs VEH = 3,820; EPI vs VEH = 383; MTX vs VEH = 1; S22-25 Tables). In line with the chromatin accessibility data, the magnitude of the response is highly correlated between ACs within each timepoint (Fig S14B), and joint modeling of the data identifies four clusters of regions (Fig S15A-B). These regions can be denoted as EAR (n = 1,079), ESR (n = 1,231), LR (n = 1,943), and NR (n = 15,026; Fig S15C and S26 Table), and show the expected change in enrichment (Fig S15D).

To determine whether the AC-responsive chromatin accessibility regions comprise regulatory regions, we overlapped our set of accessible chromatin regions (155,557) and histone-enriched regions (20,137), and compared the response to each drug (Fig 5A). Drug responses separate by treatment time, regardless of whether chromatin accessibility or active chromatin is measured (Fig S16), and within each timepoint the median drug response effect is highly correlated (3 hour, rho = 0.44; *P* < 0.001 and 24 hour, rho = 0.56; *P* < 0.001; Fig 5B). This suggests that a subset of AC-responsive chromatin changes coincide with gene regulatory regions.

**Figure 5:**
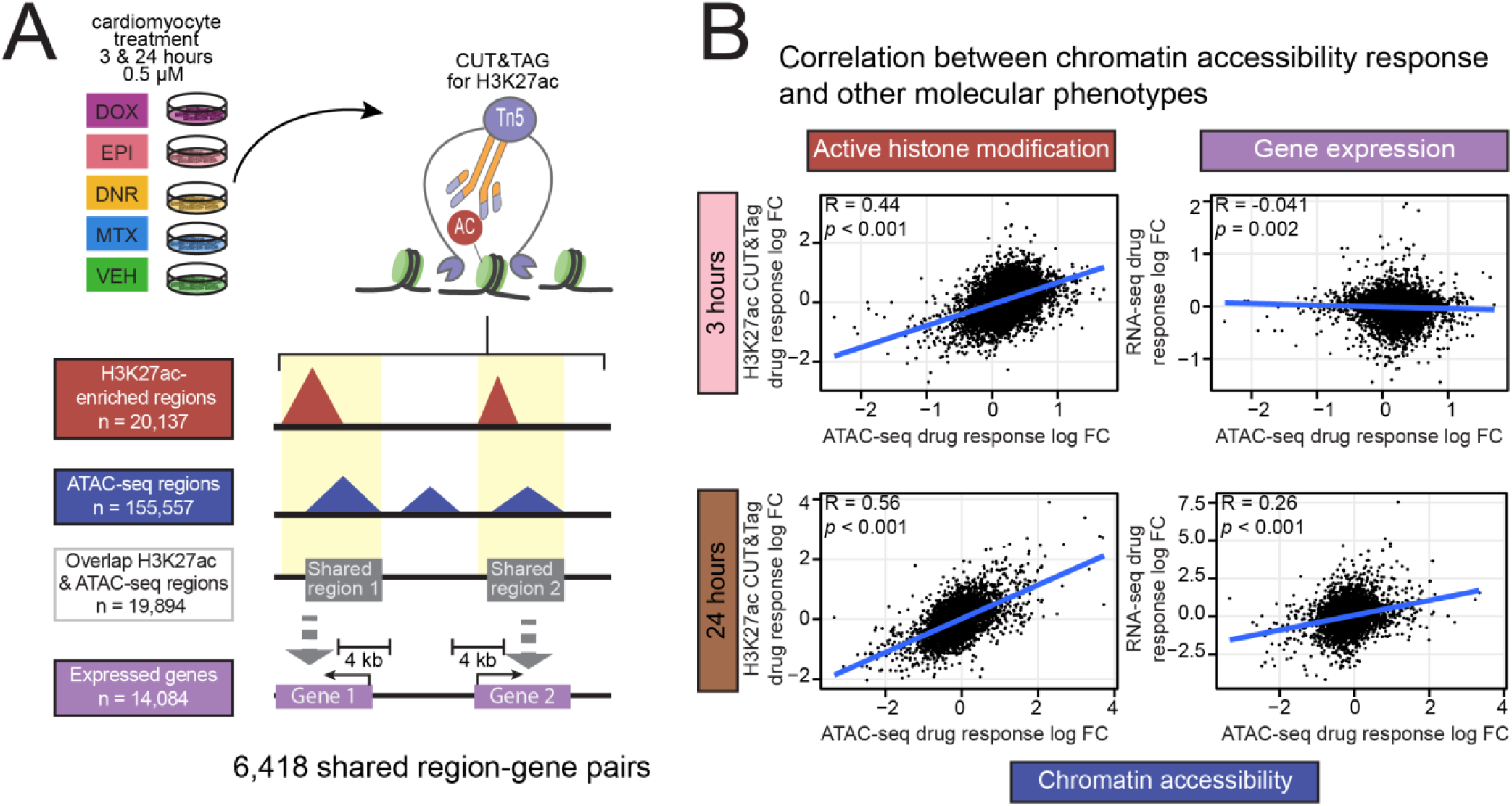
Drug-induced chromatin accessibility changes correspond to changes in active histone modification enrichment and gene expression. **(A)** Overview of the experimental set- up to collect H3K27ac CUT&Tag data and integrate with chromatin accessibility and gene expression data (15). As for ATAC-seq, iPSC-CMs were treated for three or 24 hours with 0.5 µM DOX, EPI, DNR, MTX, or VEH. H3K27ac-enriched regions are overlapped with open chromatin regions and connected to genes whose TSS is located within 4 kb of the region. **(B)** Correlation between the median log_2_ fold change (log FC) across drugs for shared region-gene pairs for chromatin accessibility and H3K27ac enrichment (left) and chromatin accessibility and gene expression (right) at each timepoint.

Considering the changes in chromatin accessibility and active histone mark enrichment in response to drug treatment, we asked whether these changes coincide with gene expression changes. To do so, we linked all accessible chromatin regions with the closest TSS within 2 kb, and correlated the response effect size (median log fold change across all TOP2i drugs) at each timepoint. We observed no correlation at three hours (rho = -0.04) and a significant correlation at 24 hours (rho = 0.26; *P* < 0.001; Fig 5B). Open chromatin regions that overlap H3K27ac show a stronger correlation with gene expression (rho = 0.26 at 24 hours; *P* < 0.001) than regions that do not overlap H3K27ac (rho = 0.15; *P* < 0.001; Fig S17). Combining drug responses across phenotypes for chromatin accessibility, active chromatin, and gene expression reveals a time-dependent effect that is shared across TOP2i (Fig S18).

### CVD-associated SNPs localize to drug-responsive chromatin regions

We next integrated our chromatin accessibility data with two GWAS that have associated AC use with CTRCD. It has been reported that there are 108 SNPs associated with AC-induced congestive heart failure (9). We asked whether any of these SNPs are associated with drug-responsive chromatin regions. Only four SNPs directly overlap our accessible chromatin regions. However, we reasoned that linked SNPs in the region may be relevant. We therefore considered accessible regions within 20 kb of the GWAS SNP. We found that 66/108 SNPs overlap with an accessible chromatin region in this window. 38/66 are associated with a drug-responsive chromatin region (Fig 6A). This includes rs10753081 and rs10798282 that fall within the same accessible chromatin region that shows increased accessibility following 24 hours of TOP2i treatment and is categorized as a late-opening response region (Fig 6B). In order to gain insight into the potential impact of the change of accessibility around this SNP, we asked whether the genotype of this SNP associates with gene expression levels in heart tissue using the GTEx portal (See Methods). Both SNPs are eQTLs for the *DARS2* gene in heart left ventricle and atrial appendage. We find that concomitant with the increase in accessibility at this SNP in response to drug treatment at 24 hours there is a decrease in expression of the *DARS2* gene (Fig 6C). Loss of *DARS2* expression leads to deregulation of mitochondrial protein synthesis (32). The closest gene to these SNPs is *PRDX6* (*DARS2* is over 350 kb away), which is not an eQTL in heart tissue but is an eQTL in seven other tissues, and its expression is also increased in response to drug treatment (Fig S19A).

**Figure 6:**
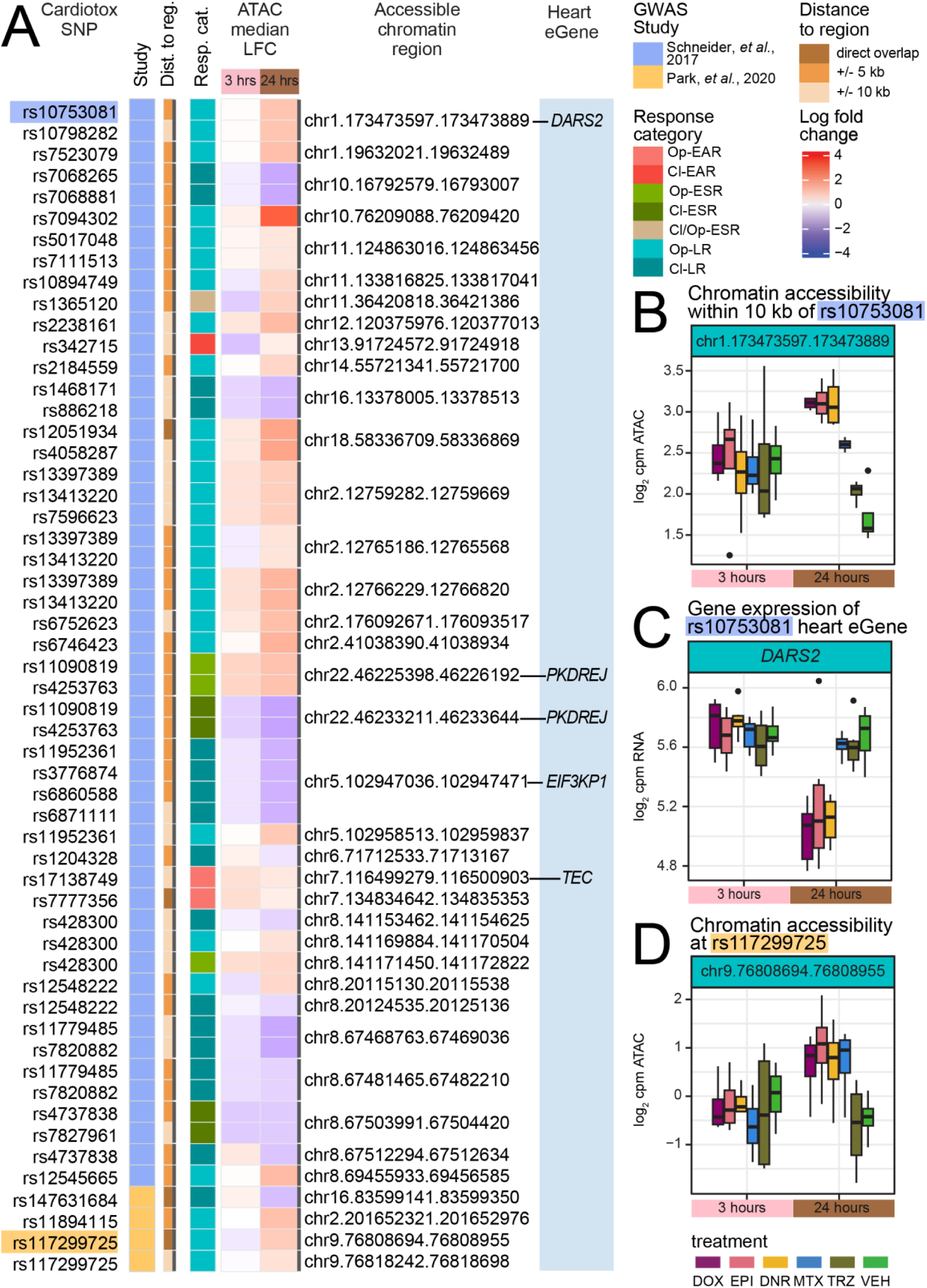
Drug-responsive regions are near cardiotoxicity-associated SNPs. **(A)** 115 AC- induced cardiotoxicity-associated SNPs were obtained from two GWAS studies (8, 9). SNPs were overlapped with drug response classes where overlaps were classified as direct (dark tan), within 10 kb (orange), or within 20 kb (light tan). The dark grey bar connects SNPs that are associated with the same chromatin region. The response class associated with the SNP is indicated as Op- EAR (pink), Cl-EAR (dark pink), Op-ESR (green), Cl-ESR (dark green), Cl/Op-ESR (beige), Op- LR (aqua blue), Cl-LR (dark aqua blue). The chromatin accessibility response to drug treatments at three and 24 hours at these loci is represented as the median log fold change across drug treatments. Cardiotoxicity-associated SNPs that are also an eQTL in heart tissue as determined by GTEx are represented as eGenes in the blue column (85). **(B)** Chromatin accessibility (log_2_ cpm) across drug treatments and time at the chromatin region that overlaps rs117299725. **(C)** Chromatin accessibility across drug treatments and time at the chromatin region within 10 kb of rs10753081. **(D)** Expression (log_2_ cpm) of the heart eGene *DARS2* (associated with rs10753081 and rs10798282) across drug treatments and time. Expression data from Matthews *et al*. (15).

A second GWAS identifies seven loci nominally associated with AC-induced cardiotoxicity (8). Four of these loci fall within 20 kb of our open chromatin regions. Two overlap directly; rs147631684, which is in a Cl-LR region in the *CDH13* gene, and rs117299725, which is in an Op-LR region in the *PRUNE2* gene (Fig 6D). This SNP is not an eQTL in any tissue, but its closest gene, *PRUNE2* shows decreased expression in response to ACs (Fig S19B).

CTRD has a range of clinical definitions, therefore we considered two types of CVD that individuals receiving ACs are at increased risk for: heart failure (HF) and atrial fibrillation (AF). We identified the set of SNPs associated with both types from the GWAS catalog (403 SNPs for HF, 679 SNPs for AF). We next overlaid the locations of these SNPs with our chromatin accessibility data. There are 32 HF SNPs and 86 AF SNPs that overlap directly with our accessible chromatin regions. 19 HF SNPs and 35 AF SNPs are in drug-responsive accessible chromatin (Fig 7A). For example, rs3176326 overlaps an Op-ESR chromatin region (Fig 7B-C), which is also an Op-LR H3K27ac region (Fig 7D). This SNP is in the first intron of the *CDKN1A* gene and is close to transcription factor motifs for ZNF460 and ZSCAN4. ZSCAN4 is known to associate with DNA damage (33). This SNP is an eQTL for *CDKN1A* in heart left ventricle. We find that as the accessibility of this region increases, the expression of *CDKN1A* increases (Fig 7E). These results indicate that genetic variants associated with CTRCD and CVD overlap regions of chromatin that change in their accessibility in response to drug treatment suggesting a mechanism behind their association.

**Figure 7:**
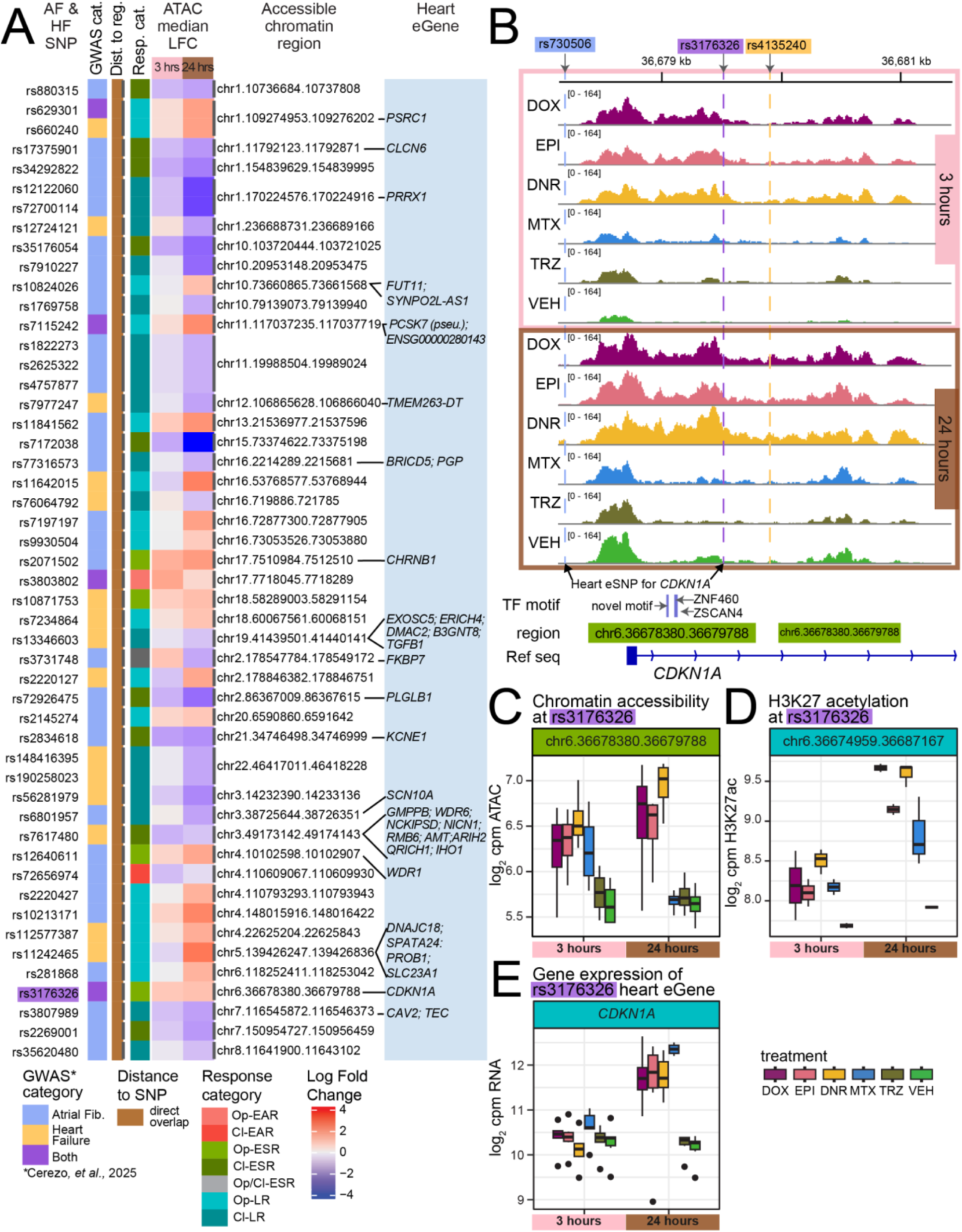
Drug-responsive regions overlap SNPs associated with atrial fibrillation and heart failure. **(A)** Heart failure (HF; gold) and atrial fibrillation (AF; light blue) associated SNPs were obtained from the GWAS catalog (11). SNPs associated with both traits are represented in purple. We overlapped the CVD SNPs with our chromatin response regions. Given the large number of SNPs (403 for HF and 679 for AF) we only considered a direct overlap (dark tan). The dark grey bar connects SNPs that are associated with the same chromatin region. The response class associated with the SNP is indicated as Op-EAR (pink), Cl-EAR (dark pink), Op-ESR (green), Cl-ESR (dark green), Cl/Op-ESR (beige), Op-LR (aqua blue), Cl-LR (dark aqua blue). The chromatin accessibility response to drug treatments at three and 24 hours at these loci is represented as the median log fold change across drug treatments. HF- or AF-associated SNPs that are also an eQTL in heart tissue as determined by GTEx are represented as eGenes in the blue column. **(B)** Chromatin accessibility at the rs3176326 SNP (purple) that is associated with both HF and AF and located within an Op-ESR chromatin response region. Accessibility from a representative individual (Individual D) is shown across drug treatments and time. Two additional CVD-associated SNPs in the region rs4135240 (HF SNP; yellow), and rs730506 (AF SNP; light blue) are also shown. Both rs730506 and rs3176326 are heart eSNPs for *CDKN1A.* Transcription factor binding motifs for ZSCAN4, ZNF460, and a novel motif were also found to be enriched within this region. **(C)** Chromatin accessibility (log_2_ cpm) across drug treatments and time at the chromatin region that overlaps rs3176326. **(D)** H3K27ac enrichment (log_2_ cpm) across drug treatments and time at the chromatin region that overlaps rs3176326. **(E)** Expression (log_2_ cpm) of the heart eGene *CDKN1A* (associated with rs3176326 and rs730506) across drug treatments and time. Expression data from Matthews *et al*. (15).

## Discussion

Individuals treated with ACs including DOX are at increased risk for developing CVD including AF and HF. DOX is a TOP2 poison, which leads to DNA double-strand breaks. The cardiotoxicity associated with DOX is mediated through interactions between the drug and the TOP2B isoform present in the heart (17, 34). In addition to DNA damage, DOX also induces damage to chromatin (35), and enhances nucleosome turnover at promoters (36). Chromatin regulators are also important contributors to survival of breast cancer patients treated with DOX (19). We previously showed that treatment of iPSC-CMs with ACs induces thousands of global changes in gene expression over time, including genes encoding chromatin regulators (15). In order to understand the contribution of chromatin to the clinical phenotype of AC-induced cardiotoxicity, and the AC-induced gene expression changes in iPSC-CMs, we profiled chromatin accessibility across five drugs used in the treatment of breast cancer. We identified and characterized tens of thousands of chromatin regions where accessibility is affected by AC treatment.

### TOP2i treatment affects the global chromatin accessibility landscape

We treated iPSC-CMs with clinically-relevant concentrations of four TOP2i and TRZ, and measured changes in accessibility at 155,557 chromatin regions three and 24 hours post-treatment. Nearly half (44%) of chromatin regions show changes in their accessibility at either the three- or 24-hour timepoint. These drug-responsive chromatin regions are shared across all TOP2i. Most changes (29%) occur 24 hours after treatment only, with few changes occurring specifically at three hours (5.2%), and 9.9% at both three and 24 hours. There are a larger number of open regions compared to closing regions at the early and late time points. The time-dependent shared drug response we observed in our chromatin accessibility data mirrors the patterns we observed in the gene expression data (15), suggesting a chromatin contribution to changes in gene expression. There is a stronger effect on chromatin accessibility than gene expression at three hours post-treatment consistent with regulatory regions becoming activated prior to the induction of gene expression changes. Indeed, we find that the magnitude of effect of chromatin accessibility changes at regions close to genes is highly correlated with the magnitude of the effect of the gene expression changes at 24 hours, but not at three hours, while the accessibility response is correlated to the change in the active histone mark H3K27ac at both timepoints. Similarly, we find chromatin response regions to be enriched for pELS identified in human heart tissue.

TRZ induces only one significant change at the three hour timepoint in chromatin accessibility in pairwise differential analysis. In the joint modeling of accessibility across drug treatments and time, there is evidence of signal in the EAR response class, albeit at a much lower probability of differential accessibility than all TOP2i treatments. There are no gene expression changes in response to TRZ at either timepoint. This suggests that TRZ induces minimal chromatin accessibility changes that result in gene expression changes, but that there are regions of chromatin that change in their accessibility soon after treatment with multiple unrelated stimuli that may or may not have an impact on gene expression.

The chromatin changes that we observe in response to ACs in iPSC-CMs are in line with the changes observed in cancer cells. DOX treatment of osteosarcoma cell lines induced ∼20,000 chromatin accessibility changes with concomitant changes in chromatin organization and condensation (37). Interestingly this study showed that etoposide, an unrelated TOP2i, had minimal effects on chromatin accessibility. We find that MTX, another unrelated TOP2i, has effects on chromatin accessibility similar to ACs. DOX treatment of retinal pigment epithelial cells induces changes in cohesin distribution leading to effects on chromatin architecture (38). Cohesin frequently binds to DNA with CTCF. We did not find an enrichment for CTCF-bound regions in heart within our drug-responsive chromatin regions; however we would need to generate CTCF or cohesin binding data in DOX-treated iPSC-CMs to determine if this mechanism is shared across cell types. Thousands of chromatin accessibility changes also exist between breast cancer cell lines that are sensitive to DOX treatment compared to those that are resistant thereby providing further support for DOX effects on chromatin (20). Interestingly, hypoxia and oxidative stress in iPSC-CMs induce minimal effects on chromatin accessibility across 15 individuals despite thousands of gene expression changes, which suggests that stress does not uniformly affect chromatin accessibility in cardiomyocytes (39).

Regions which show early increases in chromatin accessibility in response to TOP2i treatment are enriched for SVA elements. While SVAs represent a small number of TEs across the genome, they are evolutionarily young elements present only in primates, that have been shown to harbor regulatory activity. For example, they contribute to the opening of chromatin in human embryonic genome activation (40), and to species-specific *cis*-regulatory regions in liver (22, 41). Unwanted regulatory activity of SVAs is repressed in human embryonic stem cells by the ZNF91 KRAB zinc finger protein (42). Our results suggest that they may also become activated and contribute to the early response to stress. An SVA_A element that is within a chromatin region that becomes increasingly accessible over time is close to the *CAVIN1* gene, which is implicated in sensing early DNA damage (43).

Chromatin regions that increase in accessibility in response to TOP2i treatment are enriched for motifs for transcription factors implicated in the DNA damage response. This set includes p53, which is enriched in regions that are opening after 24 hours. This is consistent with p53 regulating enhancer activity by modulating chromatin accessibility (44). POU4F1, or Brn-3a, whose motif is also enriched in late response opening regions, is known to interact with p53 to enhance the activation of *CDKN1A* (45). The top ten motifs enriched in the early responsive chromatin regions bind transcription factors including NRF1, which is an adaptive stress response factor that acts as a cardioprotectant in DOX-induced cardiotoxicity (46). *NRF1* mRNA levels increase in response to DOX at both three and 24 hours further supporting its role in this system (15). Regions of chromatin that decrease in accessibility in response to TOP2i treatment are also enriched for transcription factor motifs, but do not immediately appear associated with distinct cellular pathways. We report enriched motifs as the top representative of a cluster of motifs (S14 Table). Because of the sequence degeneracy of DNA-protein binding (47), the listed motifs likely obscure a pattern of related proteins that play a role in the cardiac gene regulatory network. For example, GATA2, whose motif is enriched in Cl-EAR regions is part of the larger GATA family of transcription factors that bind the (A/T)GATA(A/G) motif (48). This includes the GATA4/5/6 family that is expressed in the developing human heart (49). The MEIS2 motif, which is enriched in Cl- LR regions is associated with defects in cardiac neural crest development (50). However, MEIS2 and MEIS1 (a transcription factor in the same motif cluster) are required for cardiac conduction and correlated with progressive cardiac conduction system disease (51). This suggests that the cardiac transcriptional landscape is impacted by closing chromatin accessibility.

### CVD-associated loci are within drug-responsive chromatin regions

Individuals treated with ACs are at increased risk for developing CVD including CTRCD, AF and HF. These CVDs have also been associated with hundreds of genetic loci in the genome. This suggests that there is both a genetic and environmental component of risk for developing CVD. Given that these loci are typically located in the non-coding genome, it can be challenging to determine the mechanistic basis for the association. Single-cell ATAC-seq experiments in human hearts have revealed that AF-associated SNPs are enriched in the cardiomyocyte population of cells suggesting that our *in vitro* system is appropriate to gain insight into disease risk (52, 53). We first asked whether loci associated with cardiotoxicity in breast cancer patients treated with DOX fall within DOX-responsive chromatin regions (8, 9). We identified a handful of variants that overlap drug-responsive chromatin regions directly, and 41 that fall within 20 kb. This includes rs117299725 that overlaps a late response opening region located in an intron of the *PRUNE2* gene directly, and rs10753081 that falls within 10 kb of a late response opening region. The latter SNP is also an eQTL for the *DARS2* gene, which we find to be downregulated in response to TOP2i.

We next investigated the overlap between our drug responsive chromatin regions and genetic loci associated with risk for AF and HF. There are hundreds of high-confidence variants associated with these traits. We identified 50 that overlap directly with drug-responsive chromatin, 21 of which overlap with opening regions. One such SNP is rs3176326 that is associated with risk for both AF and HF, and shows increased accessibility in response to TOP2i drugs. This SNP is also an eQTL for *CDKN1A*. *CDKN1A* increases its expression in response to TOP2i and is a well-characterized p53 response gene (54). These results suggest that genetic variability at these sites could affect chromatin accessibility and gene expression in response to drug treatment.

The role of chromatin modifications on risk for CVD has also been shown. H3K27ac ChIP-seq experiments in 36 failing and 34 non-failing hearts enabled the identification of histone quantitative trait loci (QTLs) (55). 22 of these loci overlap AF GWAS SNPs implying a potential regulatory mechanism for the association. Genetic variants that associate with DOX-responsive gene expression (DOX response eQTLs) have been identified in iPSC-CMs treated with DOX (14). We did not observe enrichment of DOX-response eQTLs compared to baseline eQTLs in our drug-responsive chromatin regions. This could suggest multiple gene regulatory pathways leading to DOX-responsive gene expression. Given the high degree of eQTL sharing across contexts, future work could investigate whether genetic variants associate with differences in chromatin accessibility (caQTLs). Indeed, cell-type-specific caQTLs mostly affect cell-type-specific open chromatin when comparing iPSC-CMs to the iPSCs and lymphoblastoid cells from which they were derived (56).

### Potential limitations of our study

We collected ATAC-seq data from iPSC-CMs treated with TOP2i and TRZ for three and 24 hours to determine the drug-induced chromatin accessibility landscape. We chose an early and a late timepoint; however chromatin accessibility changes may be more dynamic, and hence some effects may be absent in the timepoints we chose. While the chromatin accessibility measurements and gene expression measurements were from the same cellular differentiation, and were collected from cells treated at the same time, these were not from the same cells. More direct connections between chromatin accessibility and gene expression could be made from a single cell multi-omics study where both phenotypes are collected from the same cell. However, the strong concordance between chromatin accessibility and gene expression suggests a concerted cellular response to stress. We profiled chromatin accessibility and H3K27ac enrichment but there could be other effects on chromatin that are not captured using these approaches. Other histone modifications, transcription factors and DNA methylation may also contribute to drug-responsive gene regulation.

In summary, we have generated global chromatin accessibility data in iPSC-CMs from four individuals treated with four TOP2i drugs and TRZ, at two timepoints, to determine the impact of these drugs on chromatin. We uncovered strong effects on chromatin accessibility that associate with gene expression changes, and identified CVD-associated variants that localize in drug-responsive chromatin regions. We believe that this data from the iPSCORE panel of genotyped individuals will also provide a resource for determining the contribution of AC treatment to risk of developing CVD.

## Materials and Methods

### Ethics statement

All cell lines used were generated by Dr. Kelly A. Frazer at the University of California San Diego as part of the National Heart, Lung and Blood Institute Next Generation Consortium (57). The iPSC lines were generated with approval from the Institutional Review Boards of the University of California, San Diego and The Salk Institute (Project no. 110776ZF) and informed written consent of participants. Cell lines are available through contacting Dr. Kelly A. Frazer at the University of California San Diego, or through the biorepository at WiCell Research Institute (Madison, WI, USA).

### Induced pluripotent stem cell lines

We used iPSCs from the iPSCORE resource that were derived from four unrelated, healthy female donors of Asian ethnicity between the ages of 21 and 32 years with no previous history of cardiac disease or breast cancer (57). Individual A: UCSD143i-87-1 (iPSCORE_87_1, Asian-Chinese, age 21), Individual B: UCSD131i-77-1 (iPSCORE_77_1, Asian-Chinese, age 23), Individual C UCSD116i-71-1 (iPSCORE_71_1, Asian, age 32), and Individual D: UCSD129i-75-1 (iPSCORE_75_1, Asian-Irani, age 30).

### iPSC culture

Cells were maintained at 37 °C, 5% CO_2_ and atmospheric O_2_. iPSCs were maintained in feeder-free conditions using mTESR1 (85850, Stem Cell Technology, Vancouver, BC, Canada) with 1% Penicillin/Streptomycin (10-002-Cl, Corning, Bedford, MA, USA) on Matrigel hESC-qualified Matrix (354277, Corning, Bedford, MA, USA) at a 1:100 dilution. Cells were passaged every 3-5 days using dissociation reagent (0.5 mM EDTA, 200 mM NaCl in PBS) when the culture was ∼ 70% confluent.

### Differentiation of iPSCs into cardiomyocytes

Cardiomyocyte differentiation was performed as previously described (15). Briefly on Day 0, as a 10 cm plate of iPSCs reached 80–95% confluence, media was changed to Cardiomyocyte Differentiation Media (CDM) [500 mL RPMI 1640 (15-040-CM Corning), 10 mL B-27 minus insulin (A1895601, ThermoFisher Scientific, Waltham, MA, USA), 5 mL GlutaMAX (35050–061, ThermoFisher Scientific), and 5 mL of Penicillin/Streptomycin (100X) (30-002-Cl, Corning)] containing 1:100 dilution of Matrigel and 12 μM CHIR 99021 trihydrochloride (4953, Tocris Bioscience, Bristol, UK). Twenty-four hours later (Day 1), the media was replaced with CDM. On Day 3, after 48 hours, spent media was replaced with fresh CDM containing 2 μM Wnt-C59 (5148,

Tocris Bioscience). CDM was used to replace media on Days 5, 7, 10, and 12. Cardiomyocytes were purified through metabolic selection using Purification Media, a glucose-free, lactate-containing media [500 mL RPMI without glucose (11879, ThermoFisher Scientific), 106.5 mg L-Ascorbic acid 2-phosphate sesquimagenesium (sc228390, Santa Cruz Biotechnology, Santa Cruz, CA, USA), 3.33 mL 75 mg/mL Human Recombinant Albumin (A0237, Sigma-Aldrich, St Louis, MO, USA), 2.5 mL 1 M lactate in 1 M HEPES (L (+) Lactic acid sodium (L7022, Sigma-Aldrich), and 5 mL Penicillin/Streptomycin] on Days 14, 16 and 18. On Day 20, purified cardiomyocytes were released from the culture plate using 0.05% trypsin/0.53 mM EDTA (MT25052CI, Corning) and counted using a Countess 2 machine. A total of 400,000 cardiomyocytes were plated per well of a 12-well plate in Cardiomyocyte Maintenance Media [CMM; 500 mL DMEM without glucose (A14430-01, ThermoFisher Scientific), 50 mL FBS (MT35015CV, Corning), 990 mg Galactose (G5388, Sigma-Aldrich), 5 mL 100 mM sodium pyruvate (11360–070, ThermoFisher Scientific), 2.5 mL 1 M HEPES (H3375, Sigma-Aldrich), 5 mL 100X GlutaMAX (35050–061, ThermoFisher Scientific), and 5 mL Penicillin/Streptomycin]. iPSC-CMs were matured in culture for a further 7–10 days, with CMM Media replaced on Days 23, 25, 27, and 29.

### Differentiation efficiency determination using cardiac troponin T expression

Cardiomyocyte purity was assessed using flow cytometry and cardiac troponin T staining using the protocol described previously (15). Briefly, Day 25-27 iPSC-CMs were stained with a live stain (Zombie Violet Fixable Viability Kit (423113, BioLegend, San Diego, CA, USA)) and cardiac troponin T (TNNT2) antibody (Cardiac Troponin T, Mouse, PE, Clone: 13–11, BD Mouse Monoclonal Antibody, 564767, BD Biosciences, San Jose, CA, USA). Controls consisting of live/dead stain-only, TNNT2 antibody only, and unlabeled cells were also included. Ten thousand cells were analyzed per sample on a BD LSR Fortessa Cell Analyzer. Purity is reported as the proportion of live cells that are positive for TNNT2. Values reported are the mean of two technical replicates for each individual.

### Drug stocks and usage

The panel of drugs used were Daunorubicin (30450, Sigma-Aldrich), Doxorubicin (D1515, Sigma-Aldrich), Epirubicin (E9406, Sigma-Aldrich), Mitoxantrone (M6545, Sigma-Aldrich) and Trastuzumab (HYP9907, MedChem Express, Monmouth Junction, NJ, USA). All drugs were dissolved in molecular biology grade water to a concentration of 10 mM per stock. DOX, DNR, EPI, and MTX stocks were stored at -80 °C with working stocks stored at 4 °C for up to one week. TRZ was stored at a 1 mM concentration at 4 °C for up to one month.

### Drug treatment of iPSC-CMs and cell collection

400,000 iPSC-CMs were plated per well of a 12-well plate on Day 20 following the initiation of differentiation. On Days 27–29, iPSC-CMs were treated with 0.5 μM DNR, DOX, EPI, MTX, TRZ, or vehicle in fresh CMM. iPSC-CMs were collected three and 24 hours post-treatment, resulting in 48 samples from four individuals. iPSC-CMs were washed twice with ice-cold PBS and manually scraped using cold PBS on ice.

For the ATAC-seq samples, nuclei were extracted immediately as described below.

For the CUT&Tag samples, 100,000 manually-scraped iPSC-CMs were pelleted by centrifugation (500 g, 10 min). The supernatant was removed, and cells were resuspended in 50 µL cryopreservation-media (10% DMSO, 40% CMM, 50% FBS). Cells were placed into a CoolCell (432000, Corning) at -80 °C for 24 hours, and then stored at -80 °C until further processing.

### ATAC-seq nuclei extraction and DNA tagmentation

We processed collected iPSC-CMs using the Active Motif ATAC-Seq Kit (53150, ATAC-Seq Kit, Active Motif, Carlsbad, CA, USA) following the manufacturer’s directions. Briefly, approximately 100,000 scraped cells in PBS were pelleted by centrifugation at 500 g for 5 min. Cell pellets were resuspended in 100 μL cold ATAC Lysis Buffer. Cells were pelleted by centrifugation at 500 g for 5 min at 4 °C. Lysis buffer was removed and isolated nuclei were resuspended in 50 μL transposition mix. The tagmentation reaction was carried out at 37 °C for 30 min in a thermomixer set at 800 rpm. Transposed DNA was purified following the manufacturer’s recommendation and resuspended in 35 μL elution buffer prior to storage at -20 °C.

### ATAC-seq library preparation

ATAC-seq library preparation across the 48 samples was performed in four treatment- and time-balanced batches corresponding to all drug treatments for each individual. Libraries were prepared following the Active Motif ATAC-Seq kit manufacturer’s directions where DNA was amplified using a 14-cycle PCR reaction. Libraries were purified and bioanalyzed to determine library quality. Libraries were quantified by qPCR before being pooled and run on an Illumina Nextseq 2000 machine in four treatment- and time-balanced batches using 50 bp paired-end reads.

### CUT&Tag library preparation

CUT&Tag libraries were generated for 30 samples in three treatment- and time-balanced batches across three individuals (TRZ-treated samples were not included for this assay). Benchtop CUT&Tag was adapted from the previously described protocol (58) and Kaya-Okur & Henikoff protocols.io (https://dx.doi.org/10.17504/protocols.io.bcuhiwt6).

Cryopreserved iPSC-CMs were fast-thawed and centrifuged at 4 °C for 4 min at 1000 g to remove cryopreservation media. The pellets were resuspended and washed twice in 1 mL of Wash Buffer [20 mM HEPES-KOH pH 7.5, 150 mM NaCl, 0.5 mM Spermidine, 1x Protease inhibitor cocktail; 11873580001, Roche)]. Nuclei were extracted by incubating cells for 10 min on ice in 200 µL cold Nuclei Extraction buffer [NE Buffer; 20 mM HEPES-KOH pH 7.9, 10 mM KCl, 0.1% Triton X-100, 20% Glycerol, 0.5 mM Spermidine, 1x Protease Inhibitor cocktail]. Nuclei were pelleted by centrifugation at 4 °C for 4 min at 1,300 g. Nuclei were resuspended in PBS and fixed with 0.1% formaldehyde solution for 2 min at room temperature. Cross-linking was terminated by adding glycine to a final concentration of 0.1 M. After fixation, nuclei were centrifuged at 4 °C for 4 min at 1,300 g and resuspended in 1.5 mL of Wash Buffer.

Concanavalin A-coated magnetic beads (BP531, Bangs Laboratories Inc, Fishers, IN, USA) were prepared as previously described (59). The fixed nuclei were suspended in 1.5 mL Wash Buffer and incubated with 10 µL of activated beads for 15 min at room temperature on an end-over-end rotator. Bead-associated nuclei were collected via magnet and unbound supernatant was discarded. The sample was resuspended in 50 µL of ice-cold Antibody Buffer [Wash Buffer with 2 mM EDTA and 0.1% BSA] with a 1:50 dilution of the H3K27ac antibody (39034, Lot # 31521015, Active Motif). Samples were incubated at 4 °C overnight on a tilt table. Excess primary antibody was removed following magnetic pelleting of bead-associated nuclei. Nuclei were resuspended in 100 µL of Guinea Pig anti-Rabbit IgG (Heavy & Light Chain) preabsorbed secondary antibody (ABIN101961; Lot NE-200-032309, Antibodies-Online, Limerick, PA, USA) diluted 1:100 in Wash Buffer. The samples were incubated for 1 hr at room temperature on a tilt table. The secondary antibody was removed using a magnetic stand to clear the supernatant. The bound samples were washed twice for 5 min with 1 mL of Wash Buffer.

Tagmentation was started by incubating nuclei with 50 µL of a solution containing pAG-Tn5 (15- 1017; Lot 23243004-C1, Epicypher, Durham, NC, USA) diluted 1:20 in ‘300 Buffer’ [20 mM HEPES, pH 7.5, 300 mM NaCl, 0.5 mM Spermidine, 1x Protease inhibitor cocktail] on a tilt table for 1 hr at room temperature to bind with pAG-Tn5. Samples were collected using a magnetic stand and washed twice with 1 mL of ‘300 Buffer’. Samples were resuspended in 300 µL Tagmentation Buffer [10 mM MgCl_2_ in 300 Buffer] to activate tagmentation and incubated in a water bath at 37 °C for 1 hr. To stop tagmentation, 0.5 M EDTA, 10% SDS, and 20 mg/mL Proteinase K was added sequentially to the sample for a final concentration of 0.01 M EDTA, 0.1% SDS, and 0.15 mg/mL Proteinase K. Samples were vortexed and incubated in a water bath at 55 °C for 1 hr. DNA was extracted from the aqueous layer following PCI-Chloroform separation. The samples were chilled on ice and centrifuged at 4 °C for 10 min at 16,000 g. DNA was resuspended in 100% ethanol and centrifuged at 4 °C for 1 min at 16,000 g. DNA pellets were retained and air dried before being dissolved in 30 µL TE buffer [1 mM Tris-HCl pH 8, 0.1 mM EDTA].

Libraries were generated by combining 21 µL of DNA from each sample with 2 µL of a 10 µM uniquely barcoded i5 primer and 2 µL of a uniquely barcoded i7 primer (60). Each sample contained a unique barcode combination. A volume of 25 µL NEBNext HiFi 2 x PCR Master Mix (M0541S, New England Biolabs, Ipswich, MA, USA) was added and mixed. The sample was placed in a thermocycler with a heated lid using the following cycling conditions: 72 °C for 5 min; 98 °C for 30 s; 14 cycles of 98 °C for 10 s and 63 °C for 10 s; final extension at 72 °C for 1 min and hold at 8 °C. Post-PCR clean-up was performed by incubating libraries with 1.3 x volume of AMPureXP beads (A63881, Beckman Coulter, Brea, CA, USA) for 10 min at room temperature.

Libraries were washed twice in 80% ethanol and eluted in 25 µL 10 mM Tris pH 8.0. To eliminate very large and small DNA fragments, libraries were processed through a double-sided cleanup by adding 0.55 x the volume of AMPureXP beads followed by 1.8 x the volume of beads.

Agilent Bioanalyzer High Sensitivity DNA Analysis was performed to assess library concentrations and size distributions. Five samples failed to make libraries (Individual _A_EPI_3hour; Individual_A_EPI_24hour; Individual_B_DOX_24hour; Individual_B_MTX_3hour; Individual_C_DOX_3hour) and were removed from further processing. Libraries (n = 25) were quantified by qPCR before being pooled together. The pooled libraries were sequenced with paired-end reads and a 75 bp read length on a single lane of the Illumina NextSeq550.

### ATAC-seq analysis

#### Sequencing read processing and alignment

Raw sequencing reads were assessed for quality using FastQC (https://www.bioinformatics.babraham.ac.uk/projects/fastqc/) and visualized with MultiQC (61). Cutadapt (62) was run in PE legacy mode to remove any adapter sequences present. Paired-end sequencing reads were aligned to the hg38 genome using bowtie2 with the settings -D 20 -R 3 - N 1 -L 20 (63). Reads mapping to mitochondria and more than one genome location were removed using samtools (64). Duplicate reads were removed using Picard Tools (https://broadinstitute. github.io/picard/).

#### Identification of accessible regions

Regions of enrichment, i.e. accessible regions (peaks), were called on each file using MACS2 callpeak -f BAMPE -g hs –keep-dup all (65). A master peak set was created with BEDtools (66) by first concatenating, then merging all peaks that were adjacent (bp difference between the two peaks being considered for merging sets is 0 bp) or overlapped by at least 1 bp. We selected a set of high-confidence peaks that are present in at least five of 48 samples by first counting the total number of intersections between each .narrowPeak file with the master peak set using BEDtools multiinter function. We then removed all regions from the intersection file which had a count of < 5. Next, BEDtools intersect was used between the master peak set and the filtered intersections count file using -wa -u flags to return only those regions from the master peak file that overlapped the filtered intersection file by at least 1 bp. We removed annotated blacklist regions (https://github.com/Boyle-Lab/Blacklist/)(67) to create a master set of high-confidence open chromatin regions (n = 172,481).

#### Quantification of chromatin accessibility

To quantify the accessibility of each chromatin region in each sample we counted the number of reads in the set of high-confidence open chromatin regions using Subread featureCounts (68).

#### Fraction of reads in accessible chromatin regions

The master set of high-confidence regions (n = 172,481) was used as features in Subread featureCounts (68) to generate the fraction of paired-end reads found in these high-confidence regions. featureCounts reports the total number paired-end reads, and the number of reads that overlap at least 1 bp of the provided feature file, for each .bam file in the data set.

#### Visualization of accessible chromatin loci

The Integrative genomic viewer (IGV, version 2.19.1)(69, 70) was used to visualize open chromatin regions across the genome for all samples using the .bam files.

#### Filtering out open chromatin regions with low accessibility

The high-confidence set of 172,481 open chromatin regions were filtered to exclude regions with mean log2 cpm values < 0 and three regions mapped to the Y chromosome. This gave us a set of filtered high-confidence open chromatin regions (n = 155,557) for downstream analysis.

#### Integration with ENCODE heart ATAC-seq data

We downloaded the ATAC-seq .narrowPeak file (identifier ENCF966JZT) from ENCODE (https://www.encodeproject.org) experiment ENCSR204PZT (Michael Snyder Lab, Stanford, CA), which is derived from heart left ventricle tissue from a 41-year-old female. Using plyranges (71), we overlapped our 155,557 filtered high-confidence regions with the 218,982 regions in the .narrowPeak file.

#### TSS enrichment analysis

TSS enrichment analysis of all genes was performed on each .bam file using the TSSE function in R from the ATACseqQC package (72, 73) using hg38-known genes from the annotation package TxDb.Hsapiens.UCSC.hg38.knownGene (R package version 3.20.0). These known gene regions were used for calculating the aggregate distribution of reads centered on a TSS location over the region that flanks the corresponding TSS (TSS score) according to https://www.encodeproject.org/data-standards/terms/#enrichment. TSS score = the depth of TSS (each 100 bp window within 1000 bp each side) / the depth of end flanks (100 bp each end). TSSE score = max(mean(TSS score in each window)).

TSS enrichment of all genes within open chromatin regions was visualized using the plotAvgProf function from the ChIPseeker package in R (74, 75).

#### Identifying differentially accessible regions

To identify differentially accessible regions we used an edgeR-voom-limma pipeline (76). We first normalized the count data using TMM (Trimmed mean of M-values), then applied a voom transformation to calculate precision weights for linear modeling. Next, we modeled the individual as a random effect using the duplicateCorrelation function and transformed the data with the correlation adjustment. Finally, we contrasted each drug treatment against the vehicle at each timepoint. Differentially accessible regions (DARs) are defined as those regions for each treatment-vehicle pair that meet an adjusted *P* value threshold of < 0.05.

#### Identifying motif response classes

We jointly modeled pairs of tests to identify common accessibility patterns (or correlation motifs) that best fit the given data with a custom Cormotif R script (77). We used TMM-normalized log_2_ cpm values as input and paired each drug treatment with the corresponding VEH at each timepoint. Using the BIC and AIC, we found four motifs to be the best fit model to our data. An accessible chromatin region was considered to belong to one of the four motifs when it had a cluster-likelihood of > 0.5 of belonging to the motif and < 0.5 cluster-likelihood of belonging to any other motif. This threshold results in 98.02 % of accessible regions being assigned to an EAR, ESR, LR, or NR motif.

To understand the directionality of accessibility changes induced by each drug, we stratified each of the motifs into opening or closing regions. To define opening and closing regions, we first calculated the median log_2_ fold change (log FC) at three hours and the median log FC at 24 hours for each region across all treatments. EAR motif regions were classified as opening (Op-EAR) if the median three hour log FC > 0, and as closing (Cl-EAR) if the median three hour log FC < 0. LR motif regions were classified as opening (Op-LR) if the median 24 hour log FC > 0, and as closing (Cl-LR) if the median 24 hour log FC < 0. ESR motif regions were classified as opening (Op-ESR) if three hour and 24 hour log FC > 0, and as closing ESR (Cl-ESR) if both three hour and 24 hour log FC < 0. To capture the complexity of response, we further created an ESR opening/closing (OpCl-ESR) set defined as a median log FC at three hours > 0 and median log FC at 24 hours < 0, and an ESR closing/opening (ClOp-ESR) set defined as a median log FC at three hours < 0 and median log FC at 24 hours > 0. This stratification resulted in nine response motifs for downstream analysis.

#### Identifying TE enrichment in chromatin response regions

We obtained repeat annotations and genomic coordinates for hg38 from the RepeatMasker track(78) from the UCSC Table browser. We intersected the RepeatMasker track with our accessible chromatin regions in each response motif using the join_overlap_intersect function from the plyranges (71) package in R. We examined regions that overlapped the repeat regions by at least 1 bp. We stratified TEs by TE class: LINE, SINE, DNA, LTR, and SVA. Global TE or TE class enrichment was determined for each response motif compared to the NR motif by computing odds ratios using a 2 x 2 contingency table, and testing for significance using a chi-square test. Bonferroni adjustment for multiple testing was performed across categories of genome features. Regions with an adjusted *P* value of < 0.05 were determined to be enriched for TEs.

The SVA class was further stratified by TE family and name and proportions of each determined for the SVA-enriched accessible regions.

#### Identifying CpG island enrichment in chromatin response regions

We obtained CpG island annotations from the UCSC Table Browser (73) and overlapped these regions with our accessible chromatin regions in each response motif. We counted all regions that overlapped by at least 1 bp. Enrichment was determined by computing odds ratios using a 2 x 2 contingency table, and testing for significance using a chi-square test followed by Bonferroni adjustment for multiple testing across categories of genome features. Regions with an adjusted *P* value of < 0.05 were determined to be enriched for CpG islands.

#### Identifying TSS enrichment in chromatin response regions

We obtained TSS annotations from TxDb.Hsapiens.UCSC.hg38.knownGene package (R package version 3.20.0). and counted our open chromatin regions that overlapped these TSS by at least 1 bp for each response motif. The total number of overlapping regions in each response motif was compared to the number in the NR motif. Enrichment was determined by computing odds ratios using a 2 x 2 contingency table, and testing for significance using a chi-square test followed by Bonferroni adjustment for multiple testing across categories of genome features. Regions with an adjusted *P* value of < 0.05 were determined to be enriched for TSS.

#### Identifying heart regulatory element enrichment in chromatin response regions

We obtained a list of candidate *cis*-Regulatory Elements (cREs) from the heart-left ventricle of a female adult (46 years) from the Search Candidate cis-Regulatory Elements by ENCODE (SCREEN) database (https://screen.wenglab.org/downloads on 04/13/24). cREs were further stratified into PLS, dELS, pELS, and CTCF. We overlapped the cREs (at least 1 bp) with our accessible chromatin regions in each response motif. Enrichment was determined by computing odds ratios using a 2 x 2 contingency table, and testing for significance using a chi-square test followed by Bonferroni adjustment across categories of genome features. Regions with an adjusted *P* value of < 0.05 were determined to be enriched for cREs.

#### Identifying gene feature enrichment in chromatin response regions

We analyzed the distribution of chromatin response regions across gene features including promoter regions, exons, and introns using the annotation package TxDb.Hsapiens.UCSC.hg38.knownGene (R package version 3.20.0). Gene feature enrichment was visualized using the plotAnnoBar function from the ChIPseeker package in R (74, 75).

#### Identifying TF motif enrichment in chromatin response regions

We used the MEME Suite docker container (https://meme-suite.org v5.5.6)(79, 80) with the xstreme tool (81) that performs comprehensive motif analysis on sequences which may contain motif sites anywhere throughout the given sequences. Open chromatin regions were centered and clipped to be 200 bp +/- the center of the region for a total length of 400 bp. TF motif enrichment was determined in each chromatin response motif compared to the NR motif using the Jaspar 2022 core vertebrates non-redundant motif database to identify known motifs. The following settings were used with xstreme, --time 480 --streme-totallength 8,000,000 --meme-searchsize 100,000 --desc description --dna --evt 0.05 --minw 6 --maxw 15 --align center --meme-mod zoops --m motif_databases/JASPAR/JASPAR2022_CORE_vertebrates_non- redundant_v2.meme. Both the enrichment score and proportion of regions containing a given motif are reported.

#### Identification of expressed genes near chromatin response regions

We associated our accessible chromatin regions with the nearest expressed gene TSS using RNA-seq data for 14,084 expressed genes from these cells (15). We first obtained the TSS in the hg38 genome for all expressed genes using BiomaRt Bioconductor package in R. TSS locations were intersected with the master set of accessible chromatin regions using bedtools v2.31.0 closest function with the flags -D -b to obtain the distance of the region from the closest TSS. This created a data frame that designated the identity and distance to the nearest expressed gene for each open chromatin region used in downstream analysis.

#### Gene ontology analysis for genes near chromatin response regions

We identified the set of expressed genes with open chromatin regions within 2 kb of the TSS. A unique gene list was gathered for each response motif, and the NR motif. Gene Ontology analysis was performed with the package gProfiler2 (82, 83). The list of genes was run through the gost function with all expressed genes (n = 14,084) as a background. Significance was determined using FDR < 0.05.

### CUT&Tag sequencing analysis

#### Sequencing read processing and alignment

Raw sequencing reads for each of the 25 samples were assessed for quality using FastQC (https://www.bioinformatics.babraham.ac.uk/projects/fastqc/) and visualized with MultiQC (61). Cutadapt (62) was run in PE legacy mode to remove any adapter sequences present.

Paired-end sequencing reads were aligned to hg38 using bowtie2 with the settings –local–very-sensitive-local–no-mixed –no-discordant –phred33 -I 10 -X 700 -fr (63). Reads were filtered using samtools (64) to remove reads mapping to more than one genome location.

#### Identification of regions of H3K27ac enrichment

Enriched regions were called on each file using MACS2 callpeak -f BAMPE -g hs –keep-dup all (65). A master enriched region set was created with BEDtools (66) by first concatenating, then merging all regions that were adjacent (0 bp difference) or overlapped by at least 1 bp. We selected a set of high-confidence acetylation-enriched regions that are present in at least five of 25 samples by first counting the total number of intersections between each .narrowPeak file with the master enriched region set using BEDtools multiinter function. We then removed all intersections from the intersection file which had a count of < 5. BEDtools intersect was used between the master enriched region set and the filtered intersections file using -wa -u flags to return only those regions from the master enriched regions file that overlapped the filtered intersection file by at least 1 bp (n = 20,137).

#### Quantification of H3K27ac enrichment

To quantify the enrichment of H3K27ac in each sample we counted the number of reads in the set of high-confidence H3K27ac-enriched regions using Subread featureCounts (68).

#### Filtering out H3K27ac regions with low enrichment

The 20,137 H3K27ac-enriched regions all had mean log_2_ cpm values > 0 for each region across samples, and therefore no further filtering for low enrichment was performed.

Two samples were determined to be outliers and were removed (Fig S13, Individual_B_VEH_24hours; Individual_C_VEH_3hours).

#### Identifying differentially enriched H3K27ac regions

We performed differential enrichment analysis on H3K27ac-enriched regions. Each condition was represented by two or three individuals given the failed library and outlier samples. To identify differentially enriched H3K27ac regions, we used an edgeR-voom-limma pipeline (76). We first normalized the count data using TMM (Trimmed mean of M-values), then applied a voom transformation to calculate precision weights for linear modeling. Next, we modeled the individual as a random effect using the duplicateCorrelation function and transformed the data with the correlation adjustment. Finally, we contrasted each treatment against the vehicle at each timepoint. Differentially enriched regions are defined as those regions for each treatment-vehicle pair that meet an adjusted *P* value threshold of < 0.05.

#### Identifying H3K27ac motif response categories

We jointly modeled pairs of tests to identify common enrichment patterns that best fit the given data with a custom Cormotif R script (77). We used TMM-normalized log_2_ cpm values as input and paired each drug treatment with the corresponding VEH at each timepoint. Using the BIC and AIC, we found four motifs to be the best fit model to our data. An enriched chromatin region was considered to belong to one of the four motifs when it had a cluster-likelihood of > 0.5 of belonging to the motif and < 0.5 cluster-likelihood of belonging to any other motif. This threshold results in 95.7 % of enriched regions being assigned to an EAR, ESR, LR or NR motif.

#### Integrating ATAC-seq data with H3K27ac CUT&Tag data

155,557 ATAC open chromatin regions were overlapped with 20,137 H3K27ac-enriched regions to find shared chromatin regions using the R package plyranges (71). Shared regions are defined as regions which overlap by at least 1 bp. A total of 19,894 (98.8%) H3K27ac regions overlap open chromatin regions.

Response to drug treatment was compared across data types using the median log fold change across drug treatments at each timepoint.

#### Integrating ATAC-seq data with RNA-seq data

We associated our accessible chromatin regions with the nearest expressed gene as described above. Response to drug treatment was compared across data types using the median log fold change across drug treatments at each timepoint.

#### Identification of CVD SNPs overlapping chromatin response regions

AC-induced cardiotoxicity-associated SNPs: We obtained 108 SNPs associated with cardiotoxicity from a cohort of ∼3,000 breast cancer patients treated with an AC (9) and a list of seven SNPs associated with cardiotoxicity from a cohort of ∼3,900 breast cancer patients treated with an AC (8). The location of the SNP was obtained from the hg38 reference genome using Ensembl VEP (ensemble.org). We created windows around each SNP location corresponding to +/- 5 kb and +/- 10 kb. We overlapped SNP locations and windows with our chromatin accessibility response regions using the plyranges function join_overlap_intersect.

For all SNPs overlapping an open chromatin region, the rsID was entered into the GTEx portal (https://www.gtexportal.org/home/) to determine if the SNP is an eQTL for any gene in left ventricle heart tissue. The data were obtained on 01/30/2025.

Atrial fibrillation- and heart failure-associated SNPs: We obtained a list SNPs for atrial fibrillation (AF) and heart failure (HF) from the GWAS Catalog (https://www.ebi.ac.uk/gwas/home)(11). AF has 679 unique associated SNPs and HF 403 unique associated SNPs. Given the large number of SNPs we considered only those SNPs that directly overlap with chromatin accessibility response regions.

We determined whether SNPs overlapping open chromatin regions are an eQTL in heart tissue as described above.

## Acknowledgements

We thank all members of the Ward Lab, especially Omar Johnson, for helpful discussions. We thank Kelly Frazer and the University of California San Diego for providing the iPSC lines through the iPSCORE resource. We thank the Next Generation Sequencing Core Facility at the University of Texas Medical Branch for sequencing the ATAC-seq and CUT&Tag libraries. The Genotype-Tissue Expression (GTEx) Project was supported by the Common Fund of the Office of the Director of the National Institutes of Health, and by NCI, NHGRI, NHLBI, NIDA, NIMH, and NINDS. The data used for the analyses described in this manuscript were obtained from the GTEx Portal on 01/30/25. The authors acknowledge the Texas Advanced Computing Center (TACC) at The University of Texas at Austin for providing HPC resources that have contributed to the research results reported within this paper (http://www.tacc.utexas.edu). This work was funded by a Cancer Prevention Research Institute of Texas (CPRIT) Recruitment of First-Time Faculty Award (RR190110) to M.C.W and the National Institutes of Health grant R35GM150459 to M.C.W.

## Supporting information

S1 Appendix: Document containing

Supplemental Figures 1-19

S1 Table: ATAC-seq sample metadata

S2 Table: ATAC-seq read number and peak number by sample

S3 Table: Pairwise differential accessibility analysis for DOX vs VEH at 3 hours

S4 Table: Pairwise differential accessibility analysis for DNR vs VEH at 3 hours

S5 Table: Pairwise differential accessibility analysis for EPI vs VEH at 3 hours

S6 Table: Pairwise differential accessibility analysis for MTX vs VEH at 3 hours

S7 Table: Pairwise differential accessibility analysis for TRZ vs VEH at 3 hours

S8 Table: Pairwise differential accessibility analysis for DOX vs VEH at 24 hours

S9 Table: Pairwise differential accessibility analysis for DNR vs VEH at 24 hours

S10 Table: Pairwise differential accessibility analysis for EPI vs VEH at 24 hours

S11 Table: Pairwise differential accessibility analysis for MTX vs VEH at 24 hours

S12 Table: Pairwise differential accessibility analysis for TRZ vs VEH at 24 hours

S13 Table: ATAC-seq open chromatin regions and associated characteristics

S14 Table: Motifs and corresponding IDs found using MEME Suite for each response category

S15 Table: Significant GO:BP terms for each response category

S16 Table: H3K27ac CUT&Tag sample metadata

S17 Table: H3K27ac CUT&Tag read and peak number by sample

S18 Table: Pairwise differential acetylation analysis for DOX vs VEH at 3 hours

S19 Table: Pairwise differential acetylation analysis for DNR vs VEH at 3 hours

S20 Table: Pairwise differential acetylation analysis for EPI vs VEH at 3 hours

S21 Table: Pairwise differential acetylation analysis for MTX vs VEH at 3 hours

S22 Table: Pairwise differential acetylation analysis for DOX vs VEH at 24 hours

S23 Table: Pairwise differential acetylation analysis for DNR vs VEH at 24 hours

S24 Table: Pairwise differential acetylation analysis for EPI vs VEH at 24 hours

S25 Table: Pairwise differential acetylation analysis for MTX vs VEH at 24 hours

S26 Table: H3K27ac regions and cluster assignment

## Data availability

All ATAC-seq data have been deposited in the Gene Expression Omnibus (www.ncbi.nlm.nih.gov/geo/) under accession number GSE291260, and H3K27ac CUT&Tag data under accession number GSE291262. All custom analysis scripts used for this project are available at https://github.com/mward-lab/Matthews_cardiotox_ATAC_2025 made possible by the workflowr package (84).

## Author contributions

M.C.W conceived and designed the study. E.R.M, J.A.G, R.O.A, J.D.H performed experiments. E.R.M, R.O.A, J.D.H, M.C.W analyzed the data. E.R.M and M.C.W wrote the manuscript with input from co-authors. M.C.W supervised the work.

