## Supplementary material for "Anthracyclines induce global changes in cardiomyocyte chromatin accessibility that overlap with cardiovascular disease loci": S1 Appendix

E. Renee Matthews, Raodahtullah O. Abodunrin, John D. Hurley, Jose A. Gutierrez, Michelle C. Ward

**Table of contents**

**Figures**

Figure S1: Read numbers are similar across time and drug treatments.

Figure S2: Open chromatin region numbers are similar across time and drug treatments.

Figure S3: Samples have a high fraction of read-fragments in high-confidence open chromatin regions.

Figure S4: iPSC-CM open chromatin regions are shared with human heart-left ventricle open chromatin regions.

Figure S5: Open chromatin regions are enriched at transcription start sites.

Figure S6: Genome coverage is similar across samples at the TSS of the cardiac gene *TNNT2*.

Figure S7: ATAC-seq samples cluster by time and treatment.

Figure S8: PC1 associates with drug treatment and PC2 associates with individual.

Figure S9: Thousands of chromatin regions show changes in accessibility in response to TOP2i treatment.

Figure S10: Top differentially accessible regions are shared across anthracyclines.

Figure S11: Four differentially accessible signatures capture the response to treatment over time.

Figure S12: H3K27ac regions are enriched at transcription start sites.

Figure S13: H3K27ac CUT&Tag data separate by time and drug treatment.

Figure S14: Change in H3K27ac enrichment in response to drugs is highly correlated across ACs.

Figure S15: Four H3K27ac enrichment signatures capture the response to TOP2i over time.

Figure S16: Shared ATAC and H3K27ac region response to TOP2i clusters by time.

Figure S17: Shared accessible and H3K27ac chromatin regions have a significant correlation with nearby gene expression at 24 hours.

Figure S18: Drug response associates with time and molecular phenotype.

Figure S19: Expressed genes near SNP-containing drug-responsive chromatin regions reflect chromatin accessibility changes.

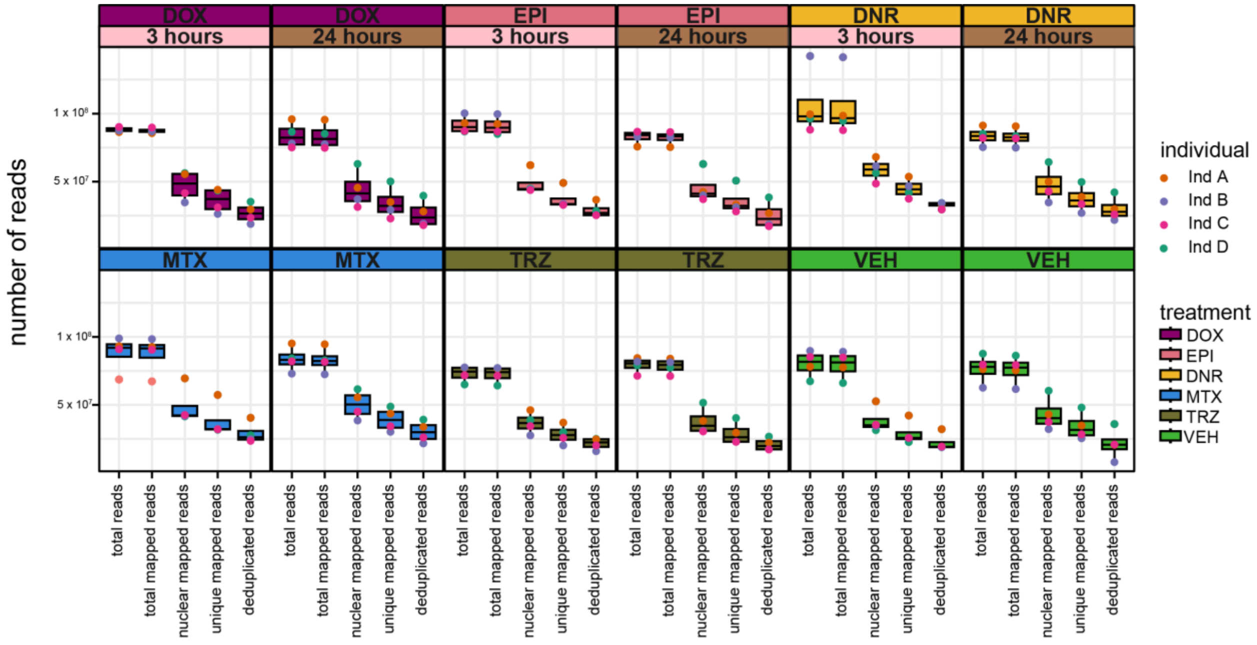


**Figure S1: Read numbers are similar across time and drug treatments.** The number of total reads, total mapped reads, total nuclear-mapped reads, unique-nuclear mapped reads, and deduplicated-unique-nuclear mapped reads by time and treatment for four individuals. Samples for each individual (A: orange; B: purple; C: magenta; D: teal) are grouped by treatment and time. Read numbers represent the sum of read 1 and read 2 for each sample.

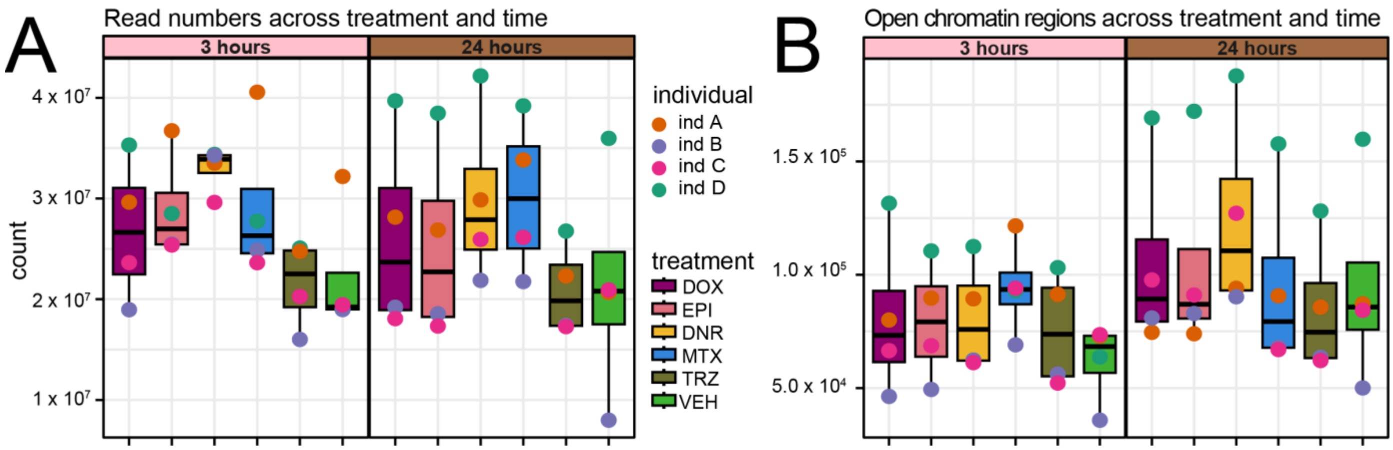


**Figure S2: Open chromatin region numbers are similar across time and drug treatments. (A)** Number of unique-deduplicated reads (read 1 + read 2 per sample) across time and treatment (DOX: mauve; EPI: pink; DNR: yellow; MTX: blue; TRZ: olive; VEH: green) at three and 24 hours for each individual (A: orange; B: purple; C: magenta; D: teal). **(B)** Number of open chromatin regions identified per treatment and time for each individual.

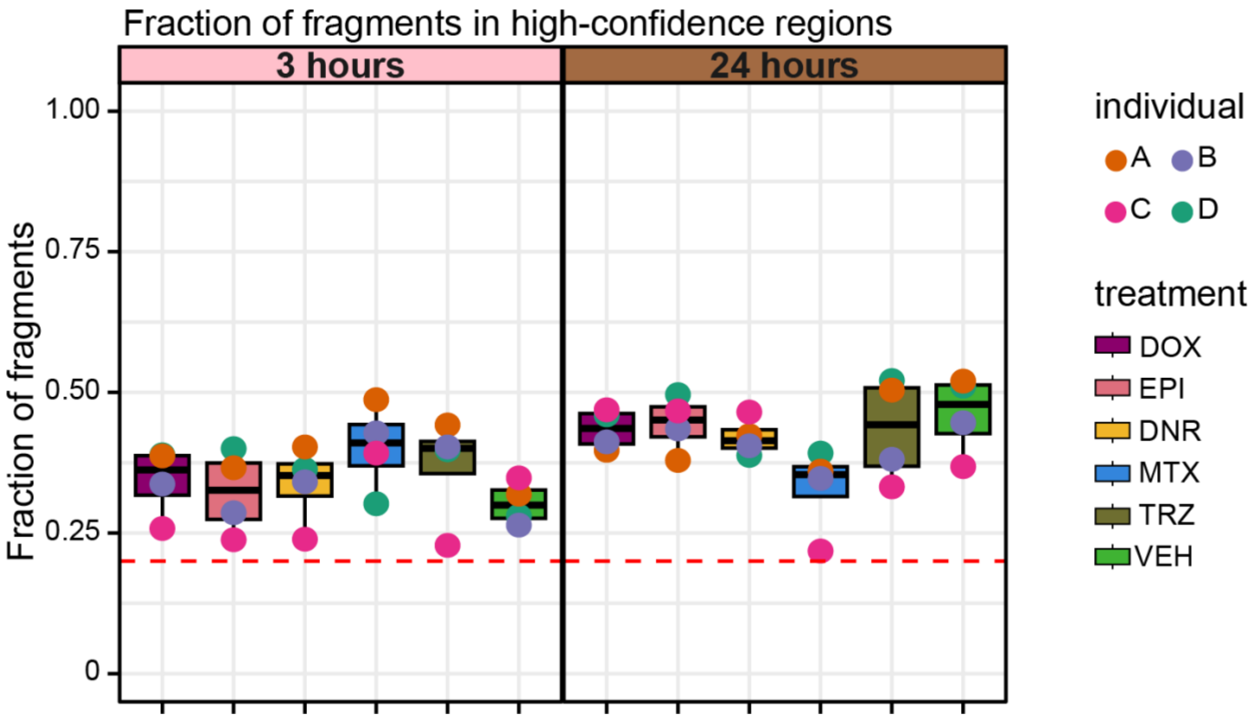


**Figure S3: Samples have a high fraction of read-fragments in high-confidence open chromatin regions.** The fraction of fragments in a set of high-confidence open chromatin regions (n = 172,481) for each individual (A: orange; B: purple; C: magenta; D: teal) across time and treatment (DOX: mauve; EPI: pink; DNR: yellow; MTX: blue; TRZ: olive; VEH: green). All samples meet or exceed the minimum acceptable quality metric of 0.2 used by ENCODE (dashed red line).

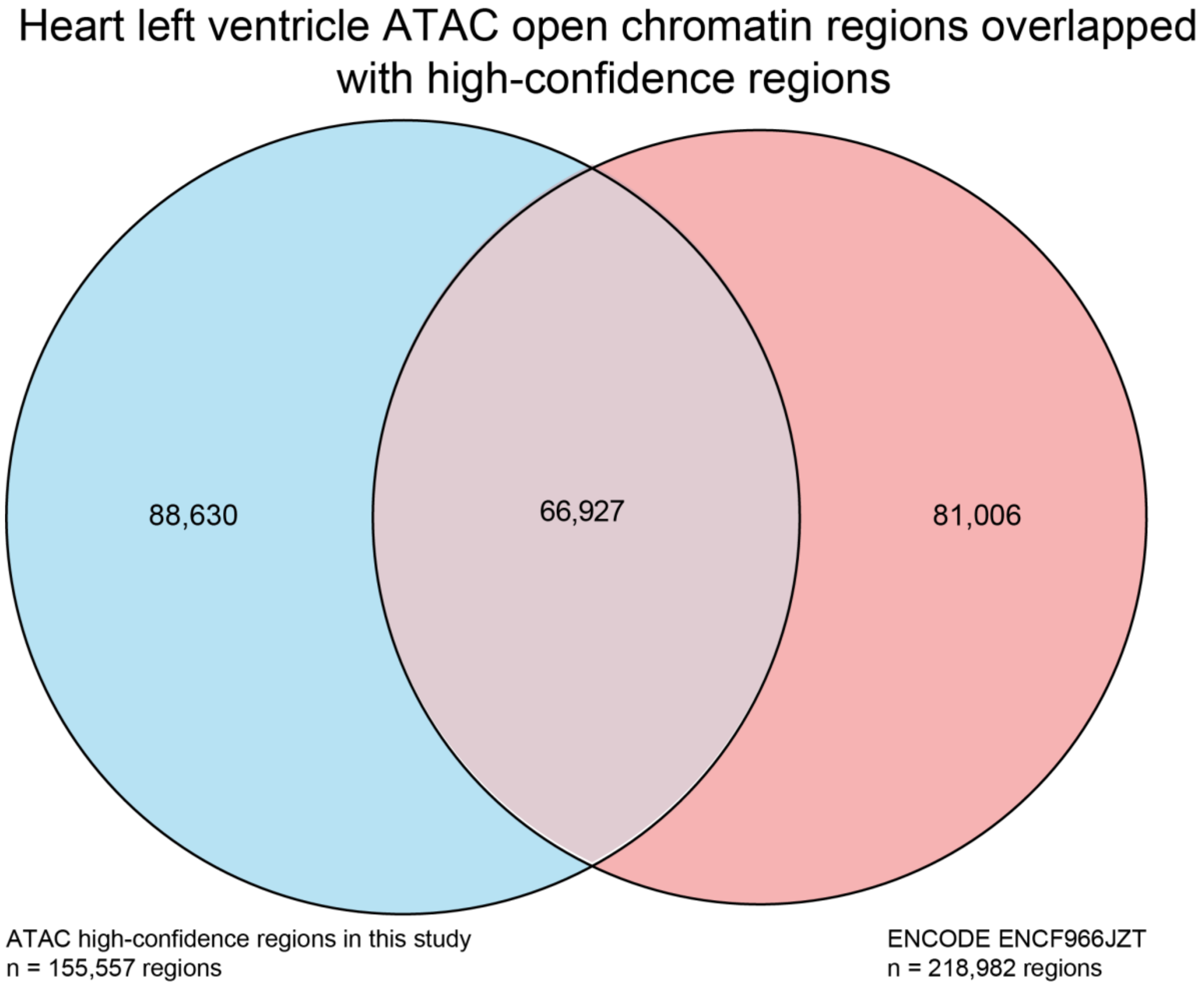


**Figure S4: iPSC-CM open chromatin regions are shared with human heart left ventricle open chromatin regions.** Overlap between our 155,557 filtered high-confidence open regions in iPSC-CMs and open chromatin regions from heart left ventricle tissue from a 41-year-old female (ATAC-seq file ENCF966JZT from ENCODE experiment ENCSR204PZT).

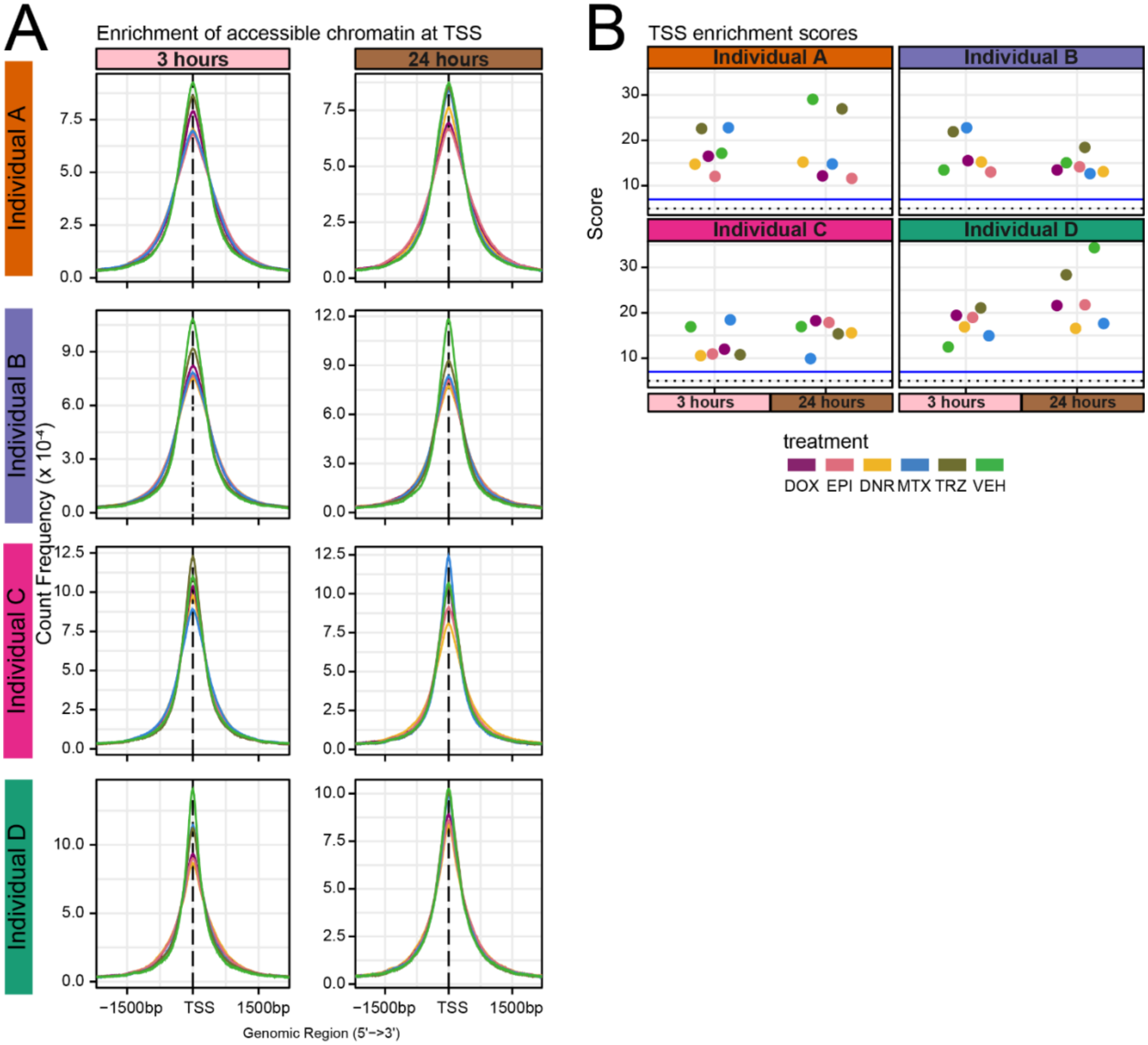


**Figure S5: Open chromatin regions are enriched at transcription start sites. (A)** Count frequency of fragments that map within +/-1.5 kb of all transcription start sites (TSS) across samples by individual and time. The dashed line is the TSS location. Solid lines are colored by treatment (DOX: mauve; EPI: pink; DNR: yellow; MTX: blue; TRZ: olive; VEH: green). **(B)** TSS enrichment scores across individuals by treatment and time. Dots represent treatment for the individuals listed above. The ENCODE ATAC-seq TSS enrichment score thresholds for the minimum ideal (solid blue line) and minimum acceptable threshold (dotted black line) are also shown.

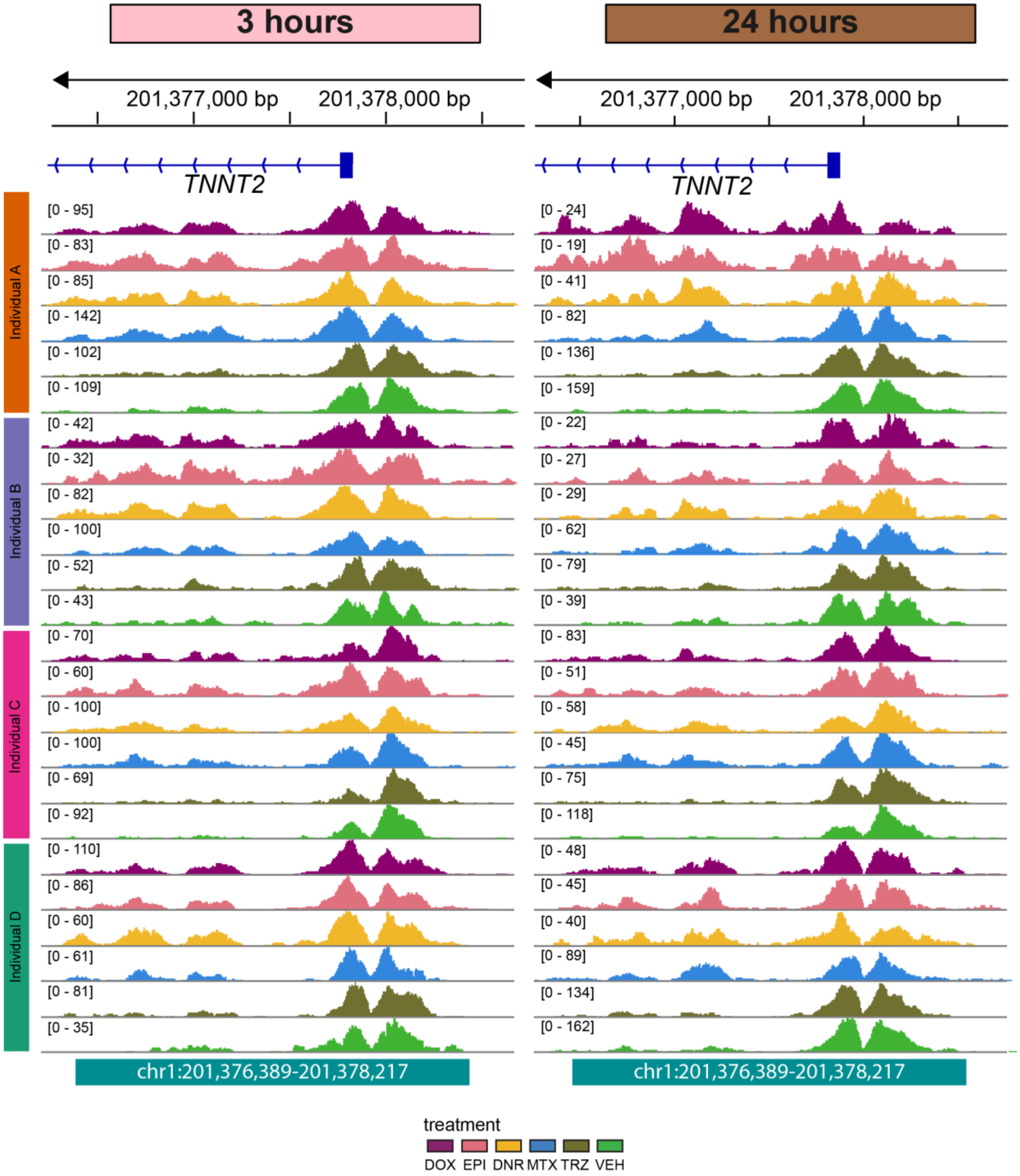


**Figure S6: Genome coverage is similar across samples at the TSS of the cardiac gene *TNNT2*.** ATAC-seq fragments at a representative open chromatin region (chr1:201,376,390-201,378,217) at the TSS of *troponin T (TNNT2)* for all samples across time and drug treatments. Three-hour samples are shown on the left, 24-hour samples are on the right. Samples are grouped by individual and colored by treatment (DOX: mauve; EPI: pink; DNR: yellow; MTX: blue; TRZ: olive; VEH: green).

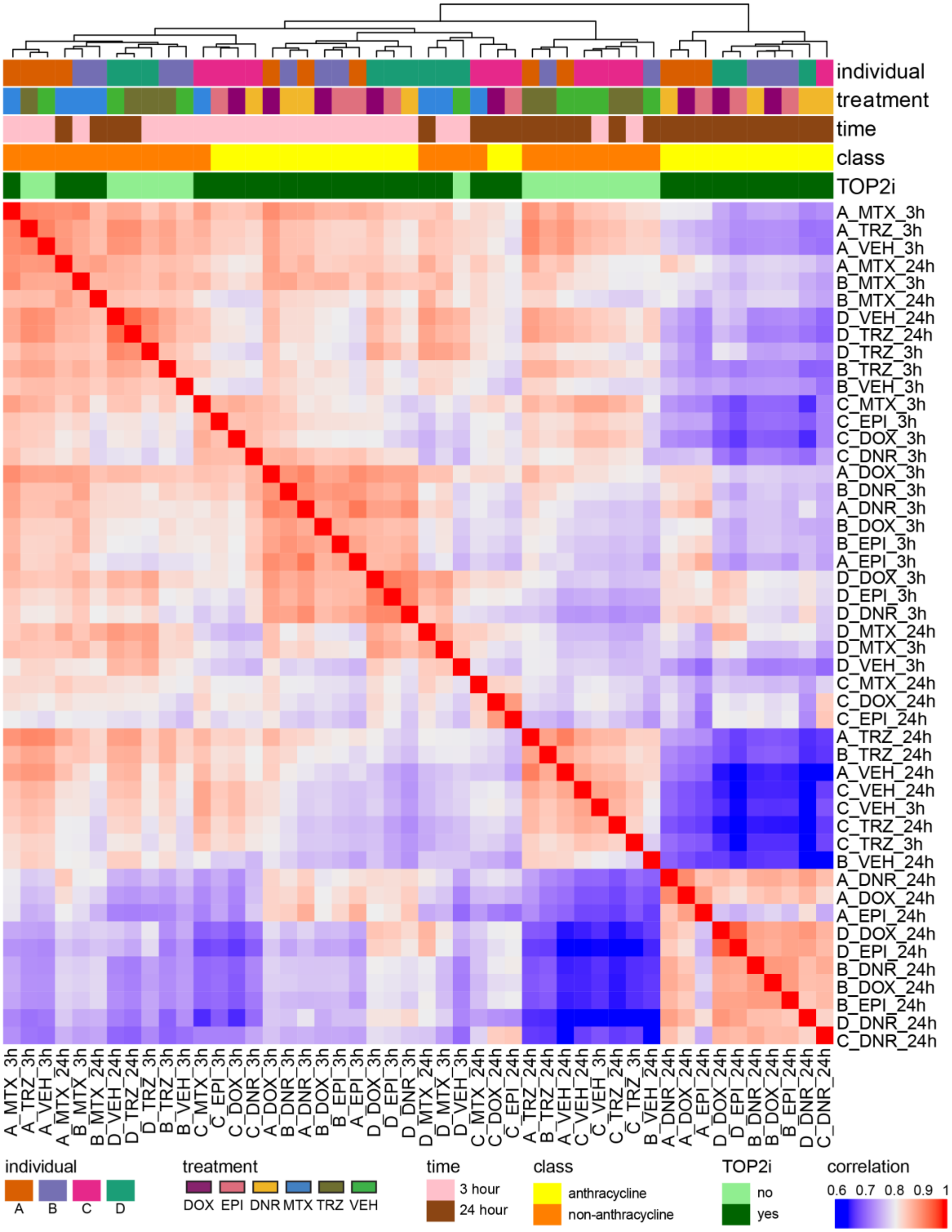


**Figure S7: ATAC-seq samples cluster by time and treatment.** Pearson correlation of $log_2$ cpm values across all pairs of samples for the high-confidence set of 155,557 open chromatin regions. Top bars are colored by individual (A: orange; B: purple; C: magenta; D: teal), treatment (DOX: mauve; EPI: pink; DNR: yellow; MTX: blue; TRZ: olive; VEH: green), time (three hours: pink; 24 hours: brown), class (anthracycline: yellow; non-anthracycline: orange), and classification as a TOP2i (no: light green; yes: dark green).

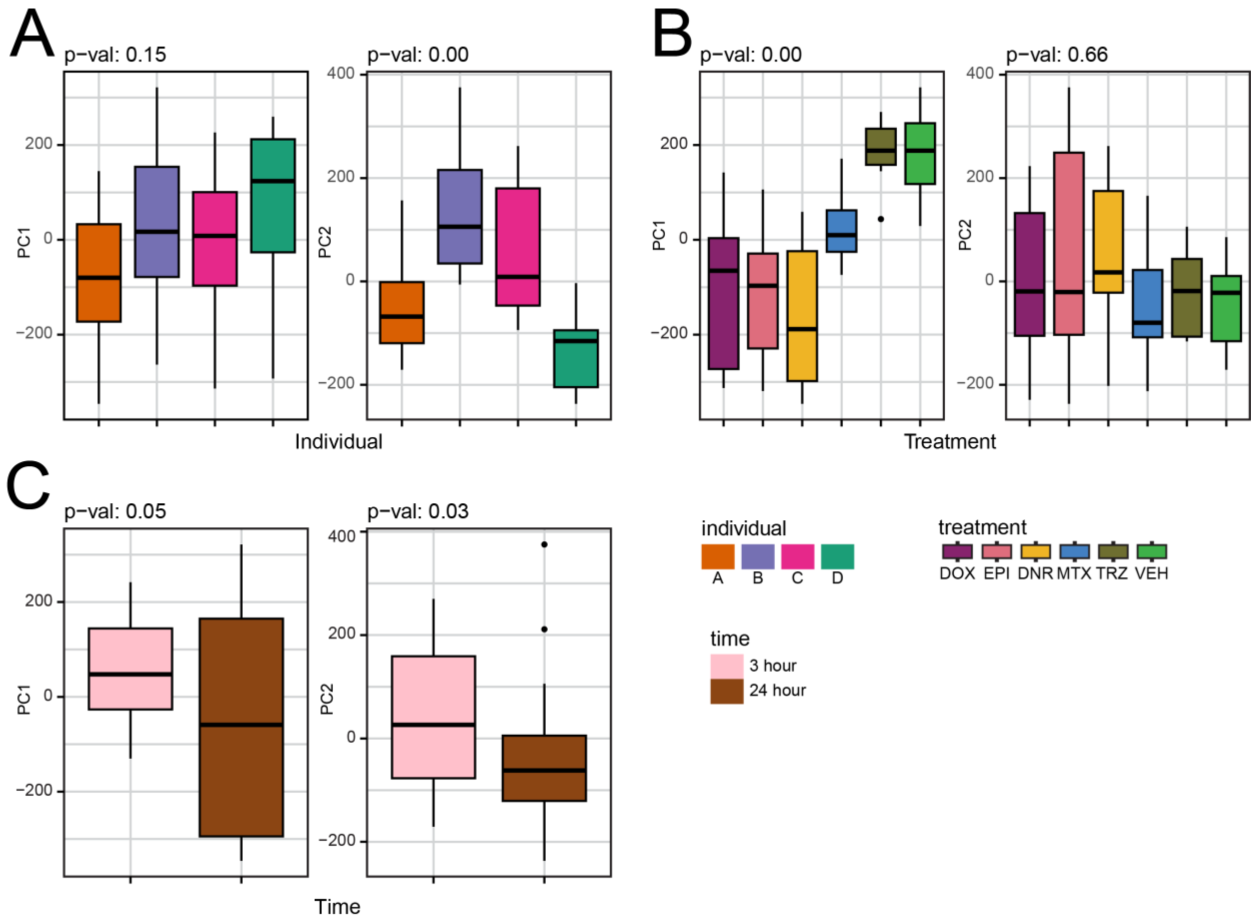


**Figure S8: PC1 associates with drug treatment and PC2 associates with individual.** Demonstration of variance contributed to the first two principal components (PC) by individual, treatment, and time (the three major biological factors in this study). **(A)** Variance of individual (A: orange; B: purple, C: magenta; D: teal) as a function of PC1 and PC2. The correlation between individual and each PC is calculated using a linear model. *P* values represent the significance of the F-statistic from the model. **(B)** Variance of treatment (DOX: mauve; EPI: pink; DNR: yellow; MTX: blue; TRZ: olive; VEH: green) as a function of PC1 and PC2. **(C)** Variance of time (three hour: pink; 24 hour: brown) as a function of PC1 and PC2.

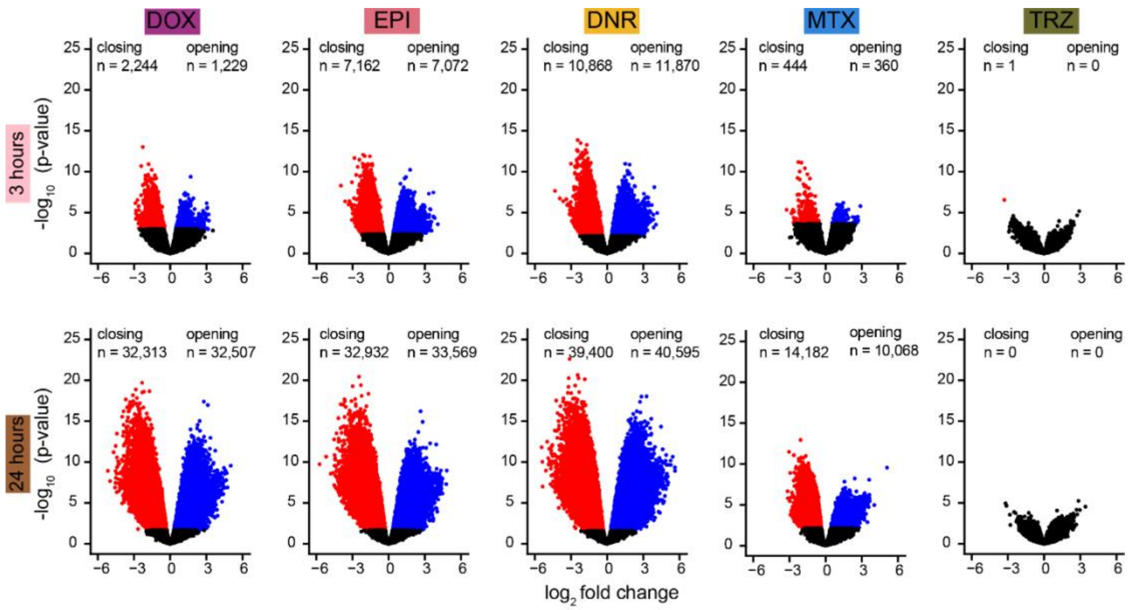


**Figure S9: Thousands of chromatin regions show changes in accessibility in response to TOP2i treatment.** Volcano plots representing open chromatin regions that are differentially accessible (adjusted *P* < 0.05) in each drug treatment compared to VEH at each timepoint. Regions that increase in accessibility in response to treatment (opening) are represented in blue, and regions that significantly decrease in accessibility (closing) are represented in red.

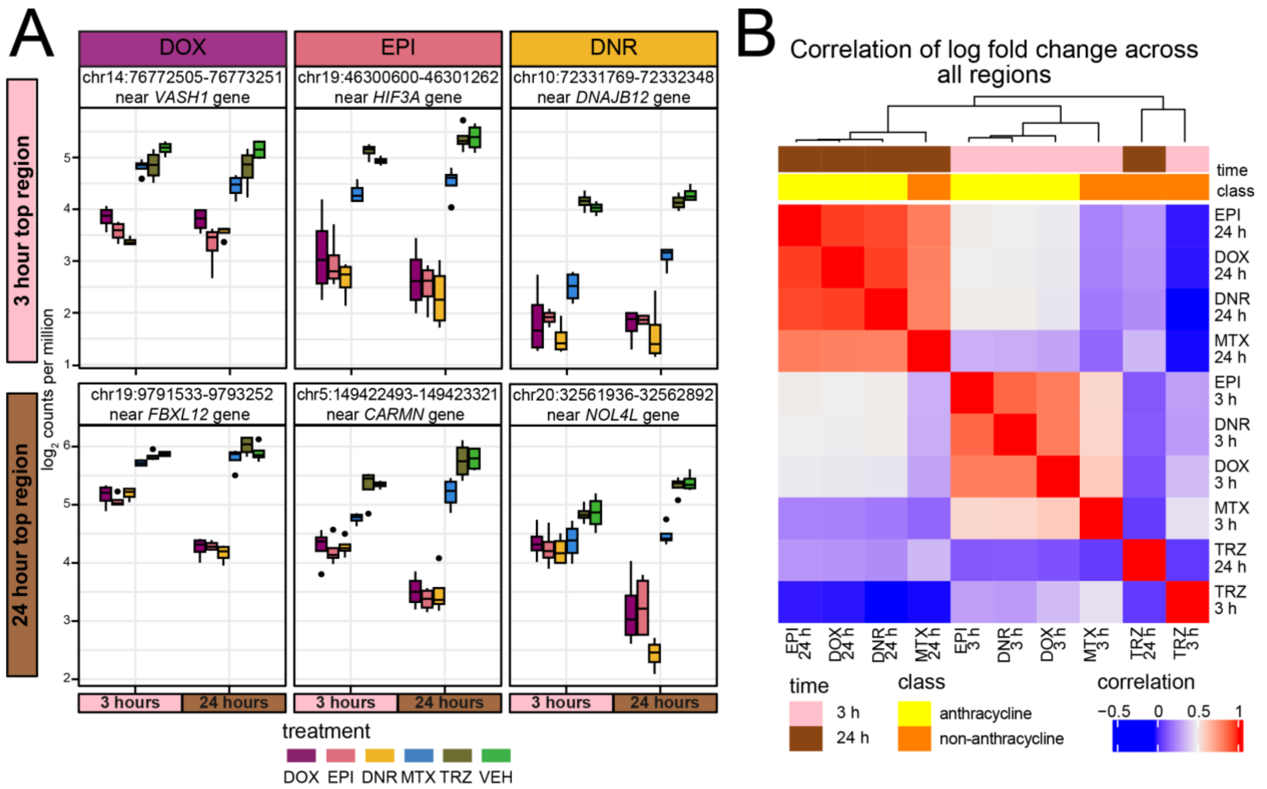


**Figure S10: Top differentially accessible regions are shared across anthracyclines. (A)** Chromatin accessibility ($log_2$cpm) at top differentially accessible regions for DOX, EPI, and DNR at three and 24 hours across each treatment (DOX: mauve; EPI: pink; DNR: yellow; MTX: blue; TRZ: olive; VEH: green). **(B)** Pearson correlation of $log_2$ fold change values across all treatment-time pairs with respect to the VEH treatment. Top color bars represent time (three hours: pink; 24 hours: brown), and class (anthracycline: yellow; non-anthracycline: orange).

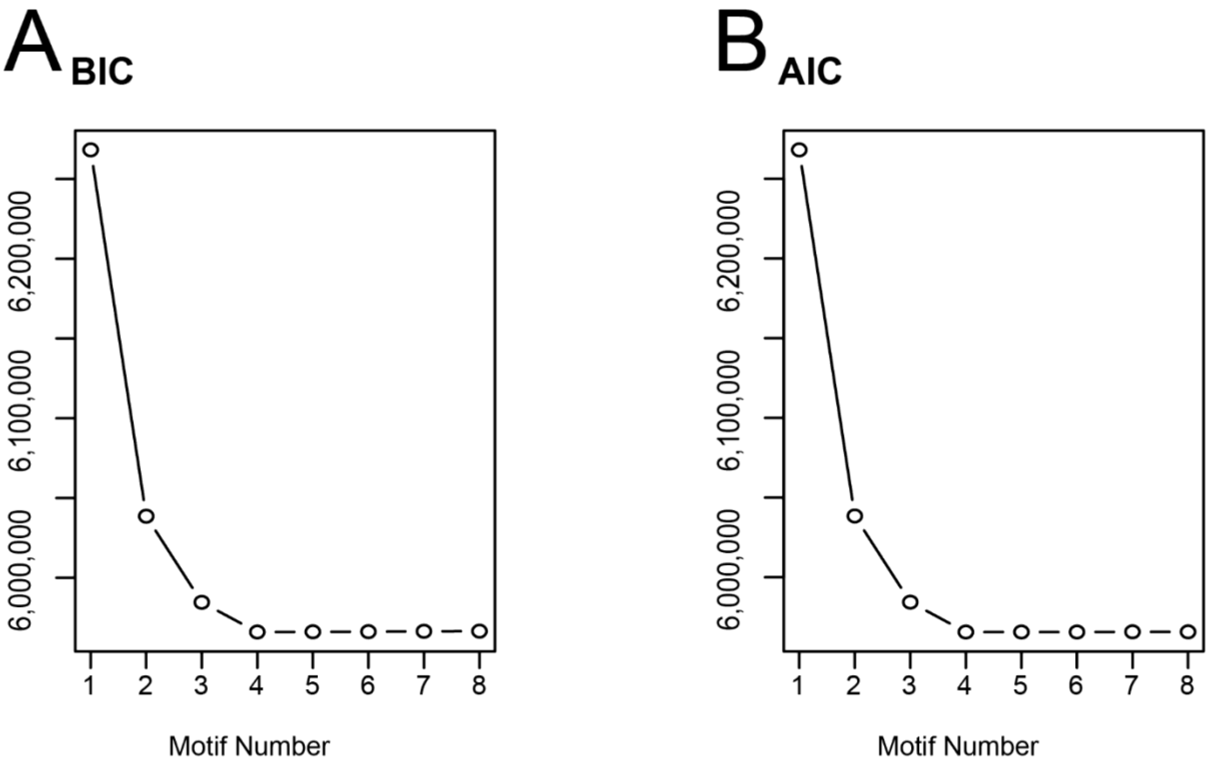


**Figure S11: Four chromatin accessibility signatures capture the response to treatment over time. (A)** Bayesian information criterion (BIC) and **(B)** Akaike information criterion (AIC) at increasing numbers of Cormotif correlation motifs following joint modeling of pairs of tests.

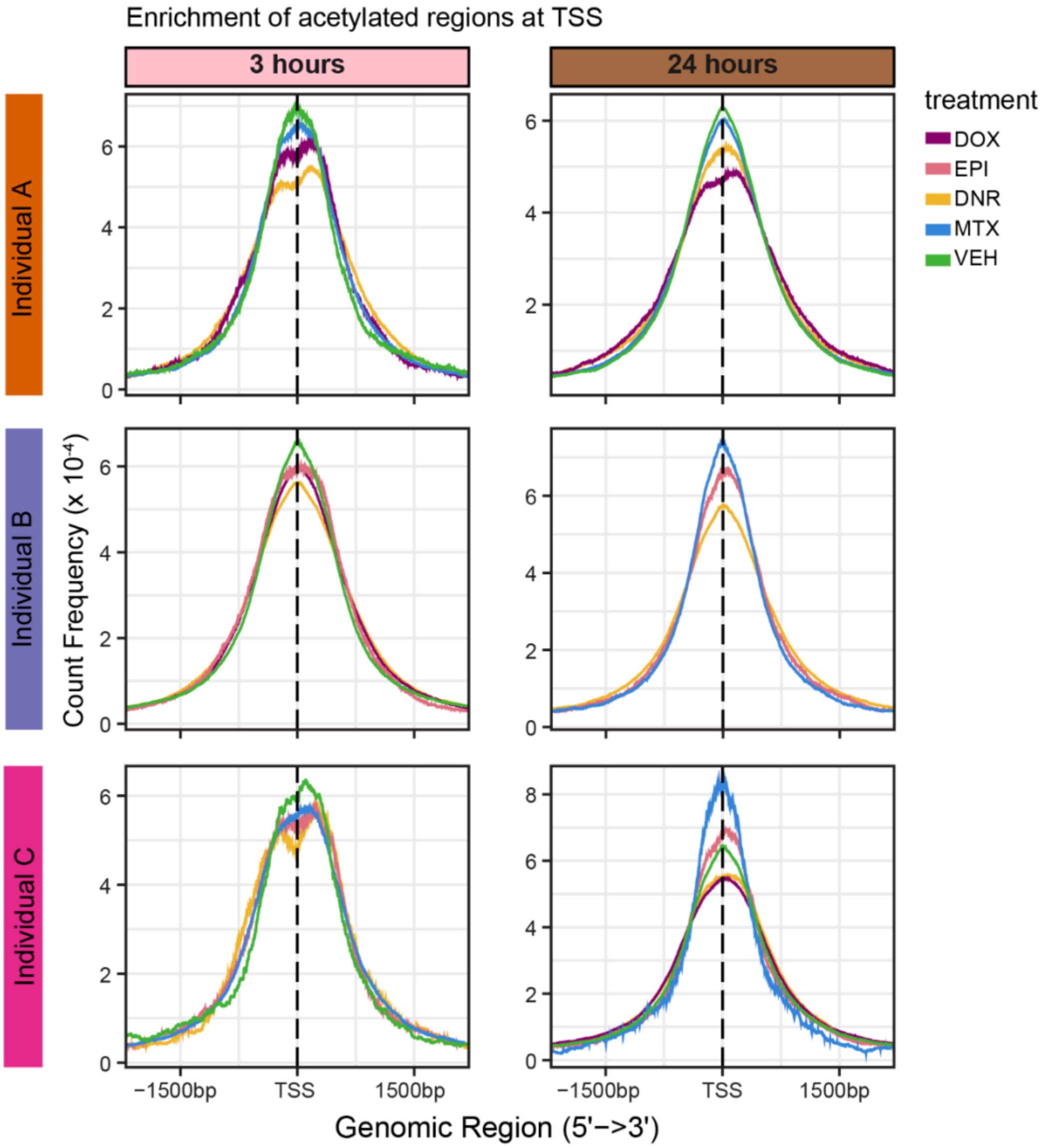


**Figure S12: H3K27ac regions are enriched at transcription start sites.** Count frequency of H3K27ac regions that map within +/-1.5 kb of all TSS in the hg38 genome across samples by individual and time. The dashed line is the TSS location. Solid lines are colored by treatment (DOX: mauve; EPI: pink; DNR: yellow; MTX: blue; VEH: light green). Individual plots with fewer than five solid lines indicate that those library preparations failed.

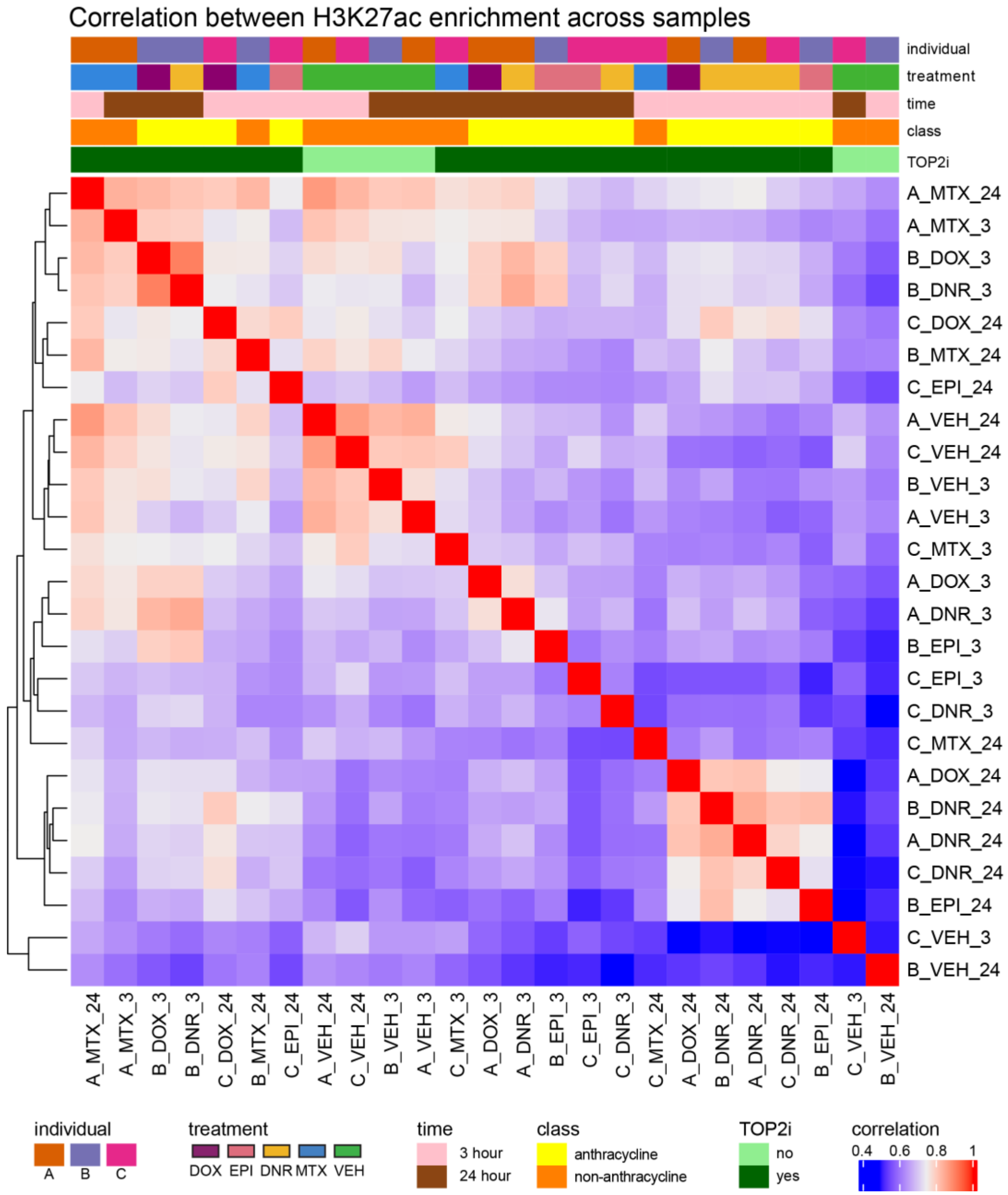


1

**Figure S13: H3K27ac CUT&Tag data separate by time and drug treatment.** Pearson correlation of $\log_2$cpm values across 20,137 high-confidence H3K27ac-enriched regions. Colored bars represent individual (A: orange; B: purple; C: magenta; D: teal), treatment (DOX: mauve; EPI: pink; DNR: yellow; MTX: blue; TRZ: olive; VEH: green), time (three hours: pink; 24 hours: brown), class (anthracycline: yellow; non-anthracycline: orange), and classification as a TOP2i (no: light green; yes: dark green).

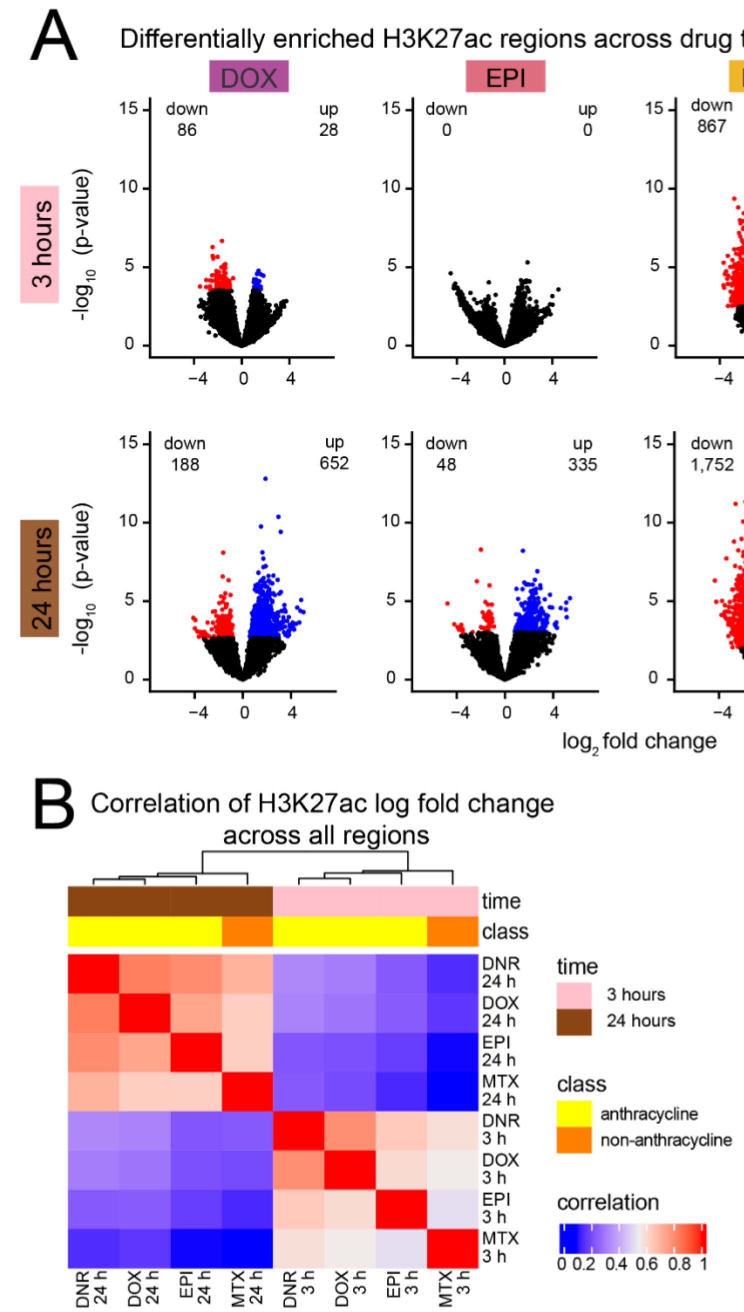


**Figure S14: Change in H3K27ac enrichment in response to drugs is highly correlated across ACs. (A)** Volcano plots representing H3K27ac enrichment changes for each drug compared to VEH at each timepoint across treatments (adjusted $P < 0.05$). Regions that increase in acetylation in response to treatment are represented in blue (up), and regions that decrease in acetylation in response to treatment are represented in red (down). **(B)** Pearson correlation of H3K27ac drug response across time and treatment. Colored bars represent time (three hours: pink; 24 hours: brown), and class (anthracycline: yellow; non-anthracycline: orange).

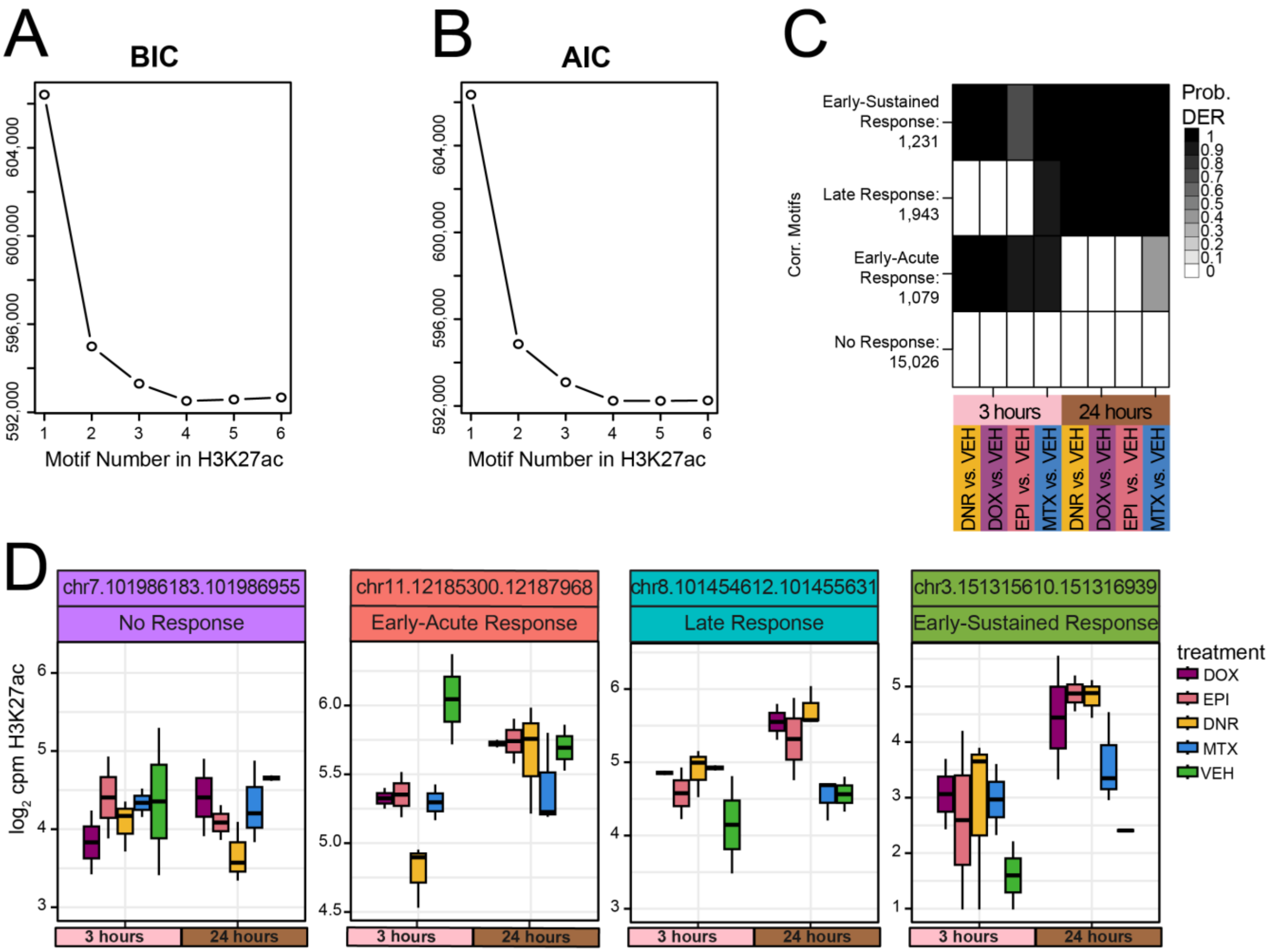


**Figure S15: Four H3K27ac enrichment signatures capture the response to TOP2i over time.** **(A)** Bayesian information criterion (BIC) and **(B)** Akaike information criterion (AIC) at increasing numbers of Cormotif correlation motifs following joint modeling of pairs of tests. **(C)** Four drug response motifs were identified following joint modeling of test pairs. Shades of black represent the posterior probability of a region being a differentially enriched region (DER) in response to the drug treatment compared to VEH at each time point. **(D)** H3K27ac enrichment ($log_2$cpm) at regions belonging to each of the H3K27ac response classes at three and 24 hours across each treatment (DOX: mauve; EPI: pink; DNR: yellow; MTX: blue; TRZ: olive; VEH: green).

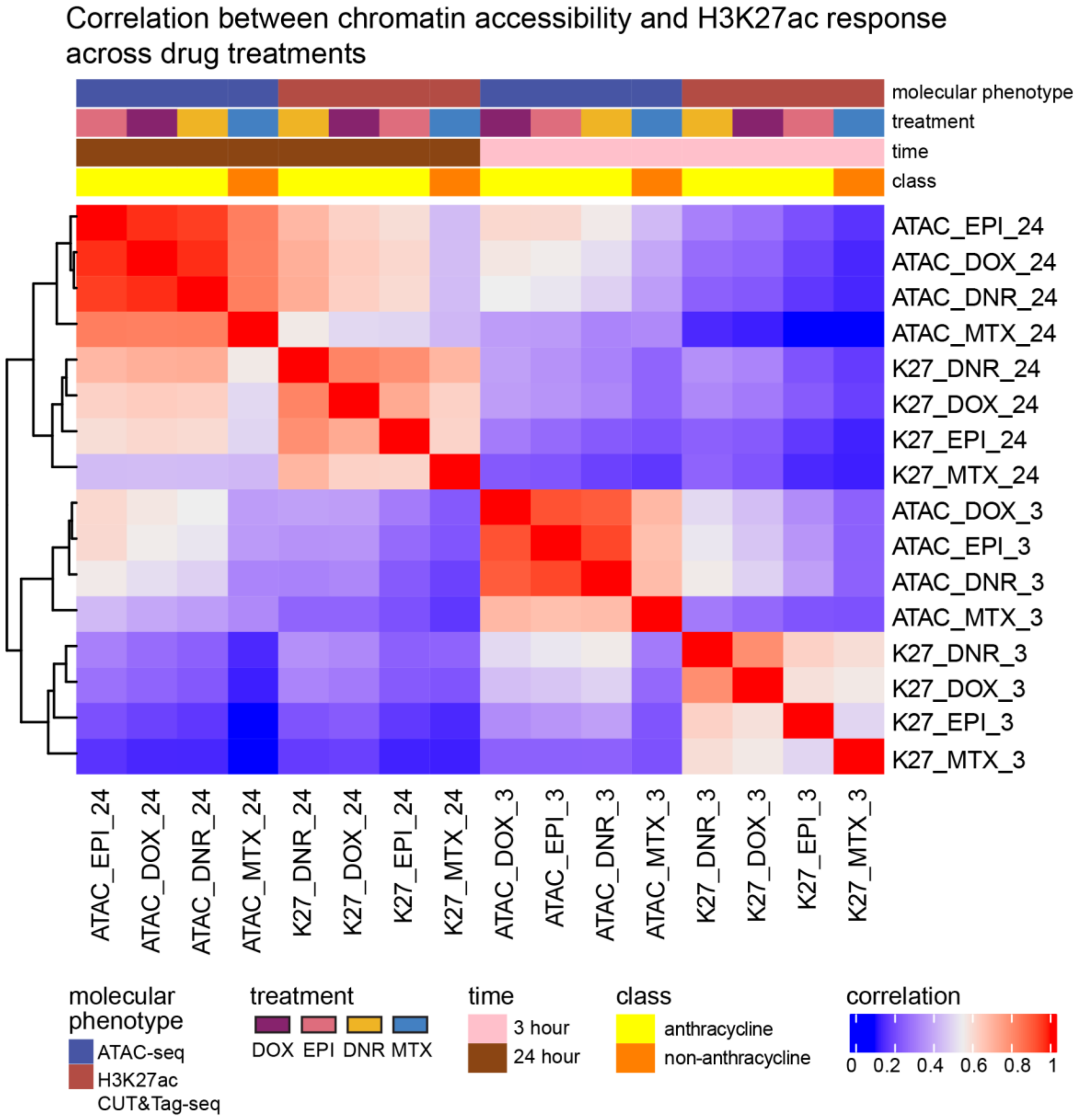


**Figure S16: Shared ATAC and H3K27ac region response to TOP2i clusters by time.** Pearson correlation of $\log_2$ fold change of drug response in regions shared between ATAC-seq and H3K27ac CUT&Tag data. Shared regions are defined as regions which overlap by at least 1 bp. A total of 19,894 (98.8 %) H3K27ac regions overlap open chromatin regions. Top bars are colored by molecular phenotype (ATAC-seq: dark blue; H3K27ac CUT&Tag: maroon), treatment (DOX: mauve; EPI: pink; DNR: yellow; MTX: blue), time (three hours: pink; 24 hours: brown), and class (anthracycline: yellow; non-anthracycline: orange).

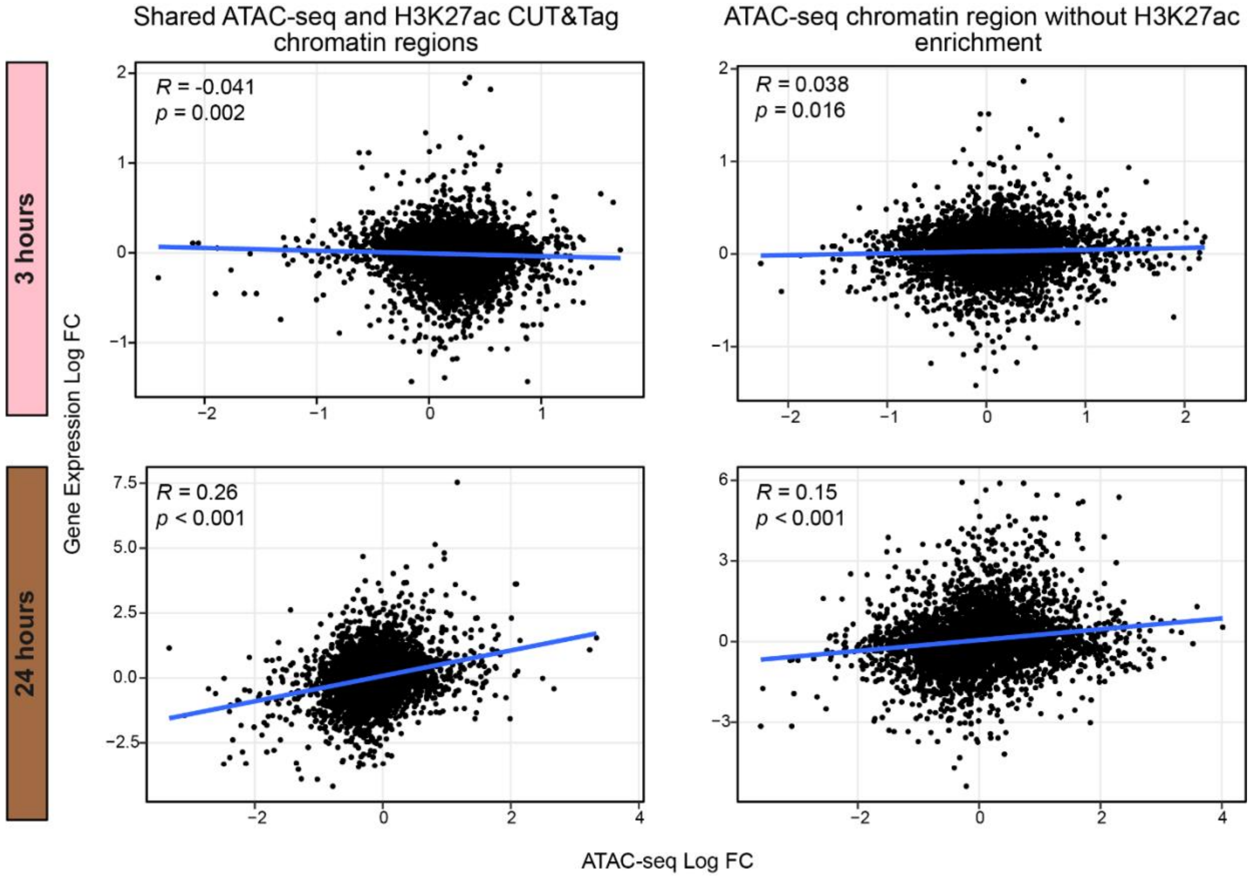


**Figure S17: Shared accessible and H3K27ac chromatin regions have a significant correlation with nearby gene expression at 24 hours.** Correlation between drug response of open chromatin regions shared with H3K27ac regions and drug response of nearby genes, and correlation between drug response of open chromatin regions that do not overlap H3K27ac regions and drug response of nearby genes. Only open chromatin regions that are within +/- 2 kb of an expressed gene TSS (Matthews *et al*., *PLOS Gen*., 2024) are shown (n = 6,418).

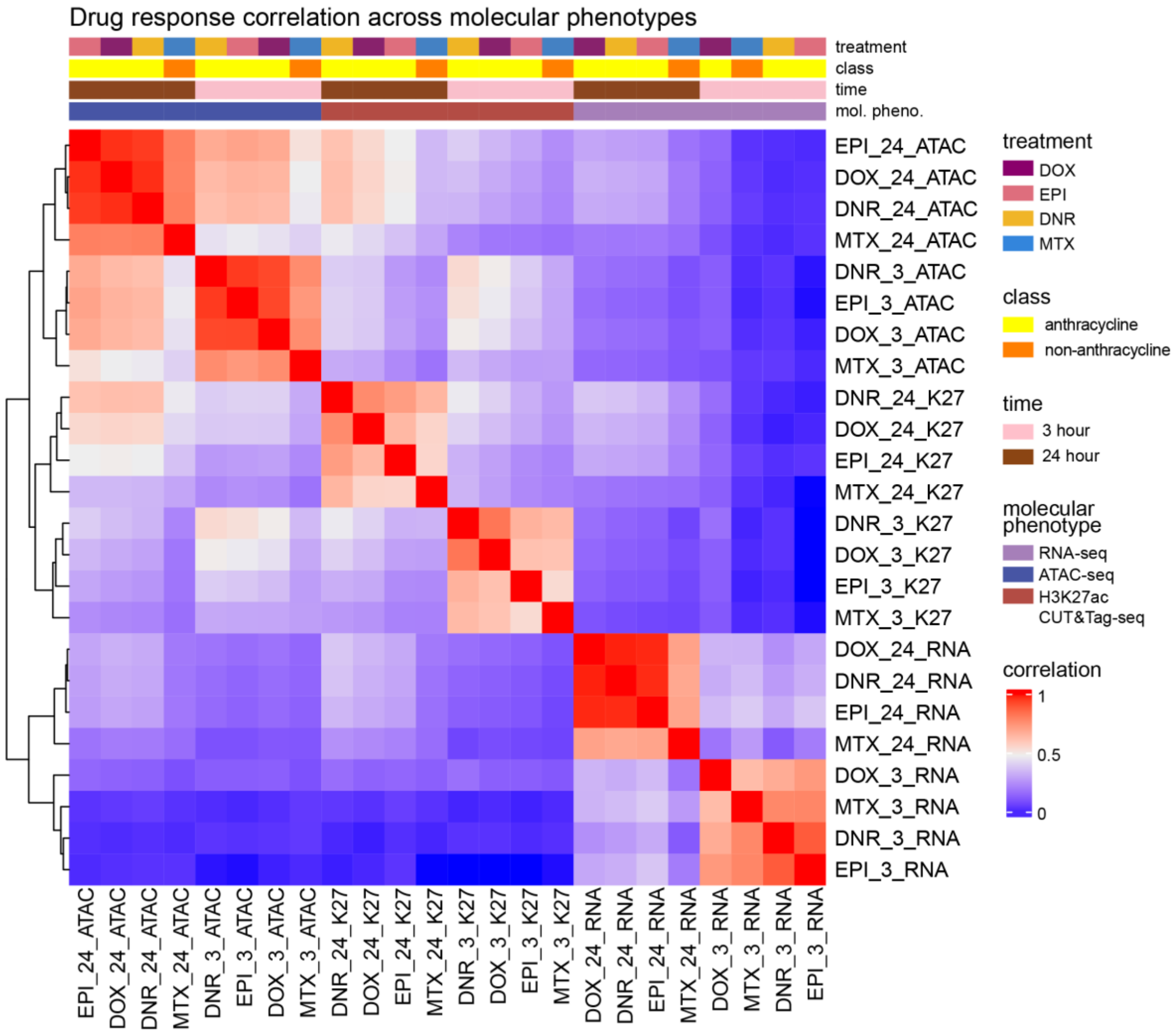


**Figure S18: Drug response associates with time and molecular phenotype.** 6,418 open chromatin regions with H3K27ac enrichment were associated with the nearest expressed gene within 2 kb of the TSS (Matthews *et al*., *PLOS Gen*., 2024). A Pearson correlation of the $\log_2$ fold change in response to each drug was performed for three molecular phenotypes (gene expression, chromatin accessibility and H3K27ac enrichment). Top bars are colored by molecular phenotype (RNA-seq: purple; ATAC-seq: dark blue; H3K27ac CUT&Tag: maroon), treatment (DOX: mauve; EPI: pink; DNR: yellow; MTX: blue), and time (three hours: pink; 24 hours: brown).

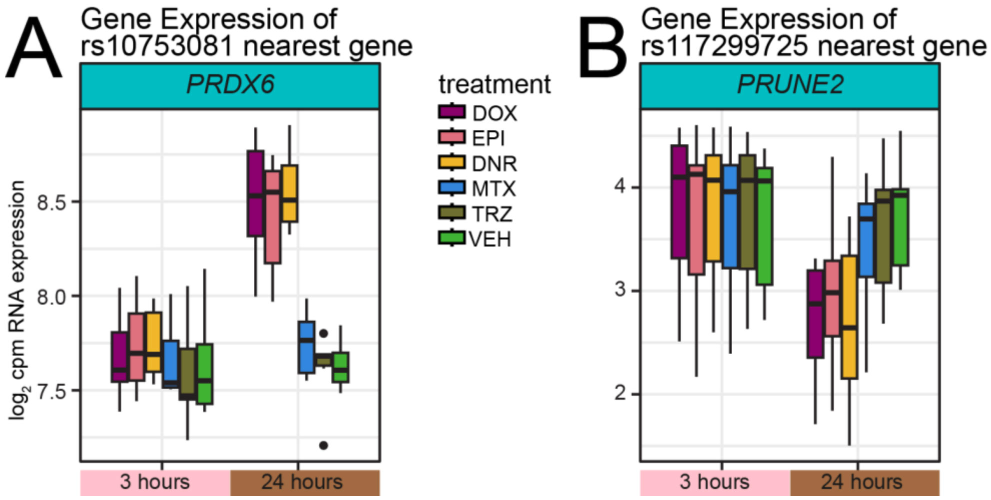


**Figure S19: Expressed genes near SNP-containing drug-responsive chromatin regions reflect chromatin accessibility changes. (A)** *PRDX6* RNA expression ($\log_2$ cpm) (Matthews *et al*., *PLOS Gen*., 2024), the closest expressed gene to the SNP rs10753081, across time and drug treatments (DOX: mauve; EPI: pink; DNR: yellow; MTX: blue; TRZ: olive; VEH: green). (**B)** *PRUNE2* RNA expression ($\log_2$ cpm)(Matthews *et al*., *PLOS Gen*., 2024), the closest expressed gene to the SNP rs117299725, across time and drug treatments.
